# Direct and Indirect Regulation of SIX1+EYA Transcriptional Activity by PA2G4, MCRS1, and SOBP

**DOI:** 10.1101/2024.09.28.615612

**Authors:** Karyn Jourdeuil, Jasmina Gafurova, Ashley Ben-Mayor, Sally A. Moody, Andre L. P. Tavares

**Affiliations:** Department of Anatomy and Cell Biology, The George Washington University School of Medicine and Health Sciences, Washington D.C., United States; Department of Biological Sciences, University of Delaware, Newark, DE, United States

**Author notes:** Contributed equally (Jasmina Gafurova, Ashley Ben-Mayor). **Corresponding Author** Dr. Andre Tavares.

**Keywords:** SIX1, EYA, Transcriptional Regulation, PA2G4, MCRS1, SOBP, BOR

## Abstract

Branchio-oto-renal (BOR) syndrome is an autosomal dominant condition characterized by variable craniofacial malformations, hearing loss and in some patients, renal dysfunction. Recently, a clinical study showed that patients with mutations causative of BOR also present with craniosynostosis, indicating that this could be an undiagnosed feature of BOR. Approximately half of all patients presenting with BOR have a variant in either *SIX1* or its activating co-factor *EYA1*. The genes underlying BOR in the other 50% of patients remain unknown. To date, most studies on the role of SIX1 in the inner ear have focused on its role in the placode-based development of the sensorineural epithelia (cochlear and vestibular organs), as sensorineural hearing loss has long term effects on human health and development. The role of SIX1 in the neural crest cells (NCCs) of the first pharyngeal (*a.k.a.*, mandibular) arch, which will give rise to the upper and lower jaws as well as the middle ear ossicles which may be affected in patients with BOR, is less well understood. To that end, we examine herein the role of three proteins that have been identified in *Xenopus laevis* as SIX1 co-factors to determine whether their function as regulators of SIX1+EYA transcriptional activity is conserved in mouse and begin to elucidate their function in the NCCs of the mandibular arch. Our data reveal that SIX1 is co-expressed with PA2G4, MCRS1, and SOBP within the oral domain of the mandibular arch, although each has a very distinct expression pattern. We further demonstrate that MCRS1 and SOBP are *bona fide* SIX1 co-factors that can repress SIX1+EYA transcriptional activity. PA2G4, on the other hand, does not bind to SIX1 in the mouse but can indirectly enhance the transcriptional activity of the SIX1+EYA transcriptional complex. Further, we show that both MCRS1 and SOBP are also able to translocate EYA to the nucleus via direct (*i.e.*, SOBP) or indirect (*i.e.*, MCRS1) mechanisms. We also show that later, during development of the mandible and incisors that SOBP is co-expressed with SIX1 within these anlagen, confirming the biological relevance of these transcriptional complexes in craniofacial bone formation and within the derivatives of the mandibular NCCs. This study, therefore, highlights the complexity of SIX1+EYA transcriptional regulation suggesting that there are multiple, and potentially redundant, mechanisms in place to ensure precise levels of SIX1+EYA transcriptional activity during craniofacial development.

## Introduction

Branchio-oto-renal (BOR) syndrome is an autosomal dominant condition characterized by variable craniofacial malformations including branchial fistulas and cysts; defects in the outer, middle, and/or inner ear that can result in hearing loss; and in some patients, renal abnormalities (Smith, 1993, Moody et al., 2015). Recently, a study by Calpena and colleagues (2022) identified that BOR-associated variants in SIX1 are present in patients presenting with craniosynostosis, suggesting that SIX1 may play an important role in suture homeostasis and that craniosynostosis may be an under-diagnosed feature of BOR. SIX1 is a homeodomain-containing transcription factor that is involved in many facets of craniofacial development (Ozaki et al., 2004, Laclef et al., 2003, Tavares et al., 2017, Guo et al., 2011, Pandur and Moody, 2000). Interestingly, SIX1 can act as either a transcriptional activator or repressor depending on the co-factors with which it interacts (Silver et al., 2003, Brugmann et al., 2004). The SIX1 protein contains a homeodomain and a SIX domain that are involved in mediating interactions with DNA and other proteins, respectively (Rafiq et al., 2021). While other members of the SIX family of transcription factors contain an activation domain (*i.e.*, SIX2, SIX4, and SIX5) and do not require the binding of a activating co-factor for transcriptional activity, SIX1 itself does not, and is transcriptionally inactive without a co-activator (Rafiq et al., 2021). The most common co-activators required for SIX1 transcriptional activity are members of the Eyes absent (EYA) family of proteins (Rafiq et al., 2021, Kenyon et al., 2005a, Kenyon et al., 2005b). EYA proteins contain a characteristic C-terminal domain, the Eya domain, which mediates their interactions with SIX proteins and other co-factors of the SIX family including the transcriptional repressor Dachshund (DACH) (Patrick et al., 2013, Kobayashi et al., 2001, Li et al., 2003, Rafiq et al., 2021, Silver et al., 2003). DACH is a highly conserved co-repressor which interacts with SIX1 to block the transcription of target genes in the absence of EYA; however, in the presence of EYA, the DACH protein acts as an activator and positively modulates the activity of SIX1 (Ikeda et al., 2002). As half of all patients with BOR present with variants in either *SIX1* or *EYA1* (Smith, 1993, Saint-Jeannet and Moody, 2014, Moody and LaMantia, 2015, Lee et al., 2007, Patrick et al., 2009, Ruf et al., 2004, Buller et al., 2001, Sanggaard et al., 2007, Li et al., 2010, Klingbeil et al., 2017), and the regulation of *SIX1* is complex and often involves multiple co-factors, we and other predict that the remaining patients for which the causative gene is unknown may have a variant in other unidentified SIX1 co-factors or downstream targets of *SIX1* (Moody et al., 2015, Neilson et al., 2010).

Many of the craniofacial structures affected in BOR are derived from the otic placode and the first (mandibular) and second pharyngeal arches (Senel et al., 2009, Kochhar et al., 2007). Early in development, neural crest cells (NCCs) migrate into the pharyngeal arches, where they receive many different cues to mediate their proliferation and differentiation into, among other things, cartilage and bone (Le Lievre, 1978, Le Lièvre and Le Douarin, 1975). Among these factors, it has been shown that the Endothelin receptor type A (EDNRA) and Bone morphogenetic protein (BMP) signaling pathways are responsible for patterning the distal-proximal axis of the mandibular arch, contributing to the organization of the lower jaw and some of the middle ear ossicles (Clouthier et al., 2000, Tavares et al., 2017, Zuniga et al., 2011, Alexander et al., 2011). The dorsal-ventral axis of the arch, on the other hand, is patterned by Jagged-Notch signaling, and loss of one copy of Jagged or Notch has been shown to result in abnormal middle ear ossicles (Barske et al., 2016, Teng et al., 2017, Zuniga et al., 2011). SIX1 is expressed along the entire length of the oral half of the mandibular arch and is required for patterning, as loss of SIX1 results in defects in the upper and lower jaw, the middle ear ossicles, and the lower incisors (Takahashi et al., 1991, Ozaki et al., 2004, Laclef et al., 2003, Tavares et al., 2017, Guo et al., 2011, Wu et al., 2019). However, as SIX1 function relies on co-factor regulation, it will be important to identify what other co-factors may be interacting with SIX1 to achieve the correct patterning of the mandibular arch.

Previous work by our group and others identified additional SIX1 co-factors that, in *Drosophila*, act as *Sine oculis* (SO, the *Drosophila* SIX1 homolog) interactors (Kenyon et al., 2005a, Kenyon et al., 2005b, Moody et al., 2015, Neilson et al., 2017, Neilson et al., 2020, Baxi et al., 2023, Jourdeuil et al., 2023, Tavares et al., 2021, Keer et al., 2022). In *Xenopus*, Proliferation associated-2G4 (Pa2G4) (Neilson et al., 2017), Microspherule protein 1 (Mcrs1) (Keer et al., 2022, Neilson et al., 2020), and Sine oculis binding protein (Sobp) (Tavares et al., 2021) function as Six1 co-factors during otic and cartilage development. Other than Mcrs1, the role of these co-factors has only been investigated during early inner ear development (Keer et al., 2022, Neilson et al., 2017, Neilson et al., 2020, Tavares et al., 2021) and their role in the NCC of the mandibular arch is largely unknown. Pa2G4, which is homologous to the fly SO interactor CG10576, contains domains thought to interact with other proteins which allow it to regulate cell proliferation and differentiation (Kowalinski et al., 2007, Monie et al., 2007, Figeac et al., 2014). In mammals, PA2G4 has been shown to interact with a wide range of proteins and its mis-regulation is associated with several types of cancers (Xia et al., 2001, Yoo et al., 2000, Lu et al., 2011, Zhang et al., 2008, Zhou et al., 2010, Kim et al., 2010, Ko et al., 2016). MCRS1 is homologous to fly dMcrs2/Rcd5/CG1135 and has been implicated in centrosome function, transcriptional regulation, and cell cycle control (Li et al., 2000, Neilson et al., 2020, Sheikh et al., 2019, Yang et al., 2015b). It is a direct binding partner of several of the proteins that make up the nonspecific lethal complex that positively influences transcription but has also been reported to modulate the transcriptional repressive activity of Death domain associated protein (Daxx), Delta-interacting protein A (DIPA), and STRA13 (Raja et al., 2010, Lin and Shih, 2002, Ivanova et al., 2005, Du et al., 2006, Shimono et al., 2005). A spontaneous recessive variant of *Sobp* causes deafness and vestibular-mediated circling behavior in the Jackson circler (*jc*) mouse (Chen et al., 2008, Calderon et al., 2006). In humans, a different *SOBP* variant has been linked to intellectual disability, anterior maxillary protrusions, strabismus, and mild hearing loss (MRAMS; OMIM #613671).

While these studies in *Xenopus* identified novel Six1 co-factors and provided functional data showing their roles during early otic and branchial cartilage development, they also demonstrated that co-factor function may change depending on the cell type into which they were transfected (*e.g.*, Pa2G4 can be a repressor or an activator of Six1+Eya1 transcriptional activity depending on the cell line used; (Neilson et al., 2017)). Therefore, it is important to confirm whether these interactions are conserved in other species and to identify the role of these putative co-factors in later craniofacial development. To this end, we examined the interactions between SIX1 and the proteins PA2G4, MCRS1, and SOBP in the mouse. EYA1 and EYA2 were also analyzed as they are required for SIX1 transcriptional activity (Li et al., 2003, Xu, 2013).

Herein, we demonstrate that MCRS1 and SOBP are *bona fide* SIX1 co-repressors in the mouse whereas PA2G4 modulates SIX1+EYA transcriptional activity without binding to SIX1. In addition, we show that SOBP can also bind to EYA and surprisingly, that PA2G4, MCRS1, and SOBP were able to translocate EYA to the nucleus without the addition of exogenous SIX1. This suggests that these co-factors may be mediating SIX1+EYA activity via both direct (*i.e.*, SOBP) and indirect (*i.e*., PA2G4 and MCRS1) mechanisms. We also demonstrate that SIX1 and its co-repressor SOBP are co-expressed during later craniofacial skeletogenesis and odontogenesis.

These findings highlight the complexity of SIX1 transcriptional regulation and show that multiple combinatorial and/or redundant mechanisms are in place to regulate craniofacial development.

## Materials and Methods

### Mice

Wild type 129S1/SylmJ mice were purchased from The Jackson Laboratory and *Six1^-/-^* (knockout) mice were kindly provided by Dr. Kiyoshi Kawakami (Jichi Medical School, Japan). All embryos used in this study were collected at embryonic day (E) 10.5 or E14.5 (Kaufman, 1994). The sex of the embryo was not determined. Genotyping of embryos from the *Six1^-/-^* mouse line was performed as described in (Ozaki et al., 2004). All experiments using mice were approved by the Institutional Animal Care and Use Committee (IACUC) at the George Washington University School of Medicine and Health Sciences (A2022-019 and A2022-020). The George Washington University Animal Research Facility is certified by the Association for Assessment and Accreditation of Laboratory Animal Care.

### Cell Transfection

Plasmids containing the full-length open reading frame (ORF) of *Mus musculus Pa2G4* (MR220724), *Mcrs1* (MR217898), and *Sobp* (MR218210) were ordered from Origene (origene.com) in the *pCMV6-Entry* Mammalian Expression Vector (PS100001). These clones were sequenced to confirm sequence homology with *Pa2G4* (NM_011119), *Mcrs1* (NM_016766), and *Sobp* (NM_175407). The ORF was subcloned into the *pCMV6-AN-Myc* Mammalian Expression Vector with an N-terminal Myc tag (PS100012). The *pCMV2-5’Flag-Six1*, *pHM6-HA-Eya1*, and *pHM6-HA-Eya2* were generously provided by Dr. Heide Ford (University of Colorado School of Medicine).

HEK293T/17 cells (ATCC CRL-11268) were cultured in Dulbecco’s modified Eagle medium (Cytivva, SH30243.01) supplemented with 10% fetal bovine serum (FBS; Gibco, 10437-028) and penicillin-streptomycin (penstrep; Gibco, 15070-063). They were plated into six-well plates (Corning, 3516) and used for co-immunoprecipitation (Co-IP) experiments because they allowed for the expression of higher levels of transfected plasmids (data not shown) and to test binding between mouse-generated proteins without interference from endogenous mouse protein intermediates. MC3T3-E1 cells (ATCC CRL-2594) were cultured in MEM α with GlutaMAX (Gibco, 32571-036) supplemented with 20% FBS and penstrep. MC3T3-E1 cells were plated into 24-well plates (Corning, 3524) for luciferase assays and into one-well Nunc Lab-Tek II Chamber slides (Thermo, 154453) for immunofluorescence (IF). MC3T3-E1 cells are a mouse pre-osteoblastic cell line with a genomic profile that closely resembles that of the cranial NC-derived mesenchyme and are therefore a good model for the NCCs contained within the mandibular arch (Wang et al., 1999, Tavares et al., 2017, Pritchard et al., 2020). Cells were transfected with Lipofectamine 3000 (Invitrogen, 2399133) for Co-IP, luciferase assays, and IF according to the manufacturer’s protocol.

### Whole mount in situ hybridization

Whole mount *in situ* hybridization (ISH) was performed as previously described (Clouthier et al., 1998). Digoxigenin-labeled probes were synthesized using the following plasmids: *pZErO-2-Six1*, *pHM6-Eya1*, *pHM6-Eya2* (kindly provided by Dr. Heide Ford), *pCRII-Pa2G4*, *pCRII-Mcrs1*, and *pCRII-Sobp* (generated by PCR and TOPO TA cloning, Thermo). To detect bound probe, E10.5 embryos were incubated in 4-nitro blue tetrazolium chloride (NBT) and 5-bromo-4-chloro-3-indolyl-phosphate (BCIP; Sigma, 5655-25TAB) except when detecting *Six1* and *Mcrs1*, for which embryos were incubated with BM-Purple (Roche, 57579200) (Tavares and Clouthier, 2015). All ISH experiments were performed on a minimum of three embryos from different litters.

### Embedding, sectioning, and immunofluorescence

E10.4 and E14.5 mouse embryos were collected in phosphate buffered saline (PBS), fixed in 4% paraformaldehyde (PFA; MP Biomedical, 150146) and processed through a graded sucrose (Amresco, 0335-1KG) series to cryoprotect tissues. All embryos were then embedded in Optimal Cutting Temperature Compound (OCT; Fisher, 4585) and stored at −80°C until sectioned. Embryo heads were serially sectioned in the transverse plane at 14 μm, collected on charged slides (FisherBrand ColorFrost, 12-550-17), and stored at −20°C until immunostained.

To immunostain sectioned E10.5 mouse embryos, slides were brought to room temperature and the tissue was rehydrated in PBS, permeabilized with 0.5% TritonX-100 (Sigma, T-9284) in PBS, and blocked for 1 hour in 0.1% TritonX-100 in PBS (0.1% PBST) with 0.5% bovine serum albumin (BSA; Sigma, A7906-100G) at room temperature. Sections were incubated in the appropriate primary antibody diluted to a 1:300 concentration in 1X PBST with 0.1% BSA (rabbit anti-EPB1 (Abcam, AB180603), rabbit anti-MCRS1 (Sigma, HPA039057-25uL), or rabbit anti-SOBP (MyBioSource.com, MBS9209963)) overnight at 4°C. Slides were washed and incubated with an appropriate secondary antibody (1:300 anti-rabbit IgG Fab2 AlexaFluor 488 secondary antibody; Cell Signaling, 4412S) overnight at 4°C. Subsequently, slides were washed with 1X PBST and incubated in a 1:300 dilution of AlexaFluor 594-conjugated rabbit anti-SIX1 (generated using Alexa Fluor 594 Antibody Labeling Kit (Thermo, A20185) and a 1 mg/mL concentrated anti-rabbit SIX1 (Cell Signaling, D5S2S)). Post-immunostaining, slides were washed with 1X PBST and cover slipped using Prolong Gold Mounting Medium with DAPI (Thermo, P36935).

For E14.5 mouse embryos, sections were serially sectioned and collected on three different slides in sequential order (i.e., slide 1A, 1B, and 1C would have transverse serial sections 14 μm apart in the same position on each slide; thus, the first section on slide 1A would be 28 μm from the same section on slide 1C but taken from the same embryo and at the same plane). The sections immunostained for rabbit anti-SIX1 and rabbit anti-SOBP were all from the same animal. For a small number of samples, a goat-anti SIX1 antibody (Santa Cruz, sc-9709) was used at a concentration of 1:500. As a control, samples were co-labeled with the rabbit-anti SIX1 from Cell Signaling and the goat-anti SIX1 to ensure that the labeling was identical in these samples (data not shown). All immunostaining was performed as described for E10.5 above. All immunostaining of mouse embryos was performed in triplicate.

Immunostaining was also performed on MC3T3-E1 cells transfected with different combinations of *pCMV6-AN-Myc* (control), *pCMV2-5’Flag-Six1*, *pHM6-HA-Eya1*, *pHM6-HA-Eya2*, *pCMV6-AN-Myc-Pa2G4*, *pCMV6-AN-Myc-Mcrs1*, and/or *pCMV6-AN-Myc-Sobp*. 1.5 μg of each plasmid was transfected into cells plated on one-well Nunc Lab-Tek II Chamber slides. Cells were processed as previously described (Shah et al., 2020). Briefly, 48 hours post transfection, cells were fixed in 4% PFA for 10 minutes and processed for immunostaining by standard methods using combinations of rabbit (Cell Signaling, 71D10) or mouse anti-Myc (Cell Signaling, 9B11), rabbit (Cell Signaling, C29F3) or mouse anti-HA (Cell Signaling, 6E2), and/or rabbit anti-SIX1. Slides were then incubated with anti-rabbit Alexa Fluor 488 and anti-mouse IgG (H+L) Fab2 Fragments Alexa Fluor 594 (Cell Signaling, 8890S), washed, and cover slipped with Prolong Gold Mounting Medium with DAPI. Experiments were repeated at least three independent times and at least five fields per slide were analyzed.

Sections were imaged using a Zeiss LSM710 or LSM910 confocal microscope (high magnification images). Low magnification images were acquired with either a Leica DM6000B or a Leica Observer.Z1 upright microscope. Transfected MC3T3-E1 cells were imaged using only the LSM710 confocal microscope. Images were processed using Adobe Photoshop 2022 or Adobe Photoshop 2024. The protein-specific antibodies for SIX1, PA2G4, MCRS1, and SOBP were validated via Western blot to bind specifically to the protein of interest using lysate from MC3T3-E1 cells transfected with the tagged proteins of interest (see **Supplemental Fig. 1**).

### Co-Immunoprecipitation

HEK293T/17 cells were transfected with combinations of *pCMV6-AN-Myc-control*, *pCMV2-5’Flag-Six1*, *pHM6-HA-Eya1*, *pHM6-HA-Eya2*, *pCMV6-AN-Myc-Pa2G4*, *pCMV6-AN-Myc-Mcrs1*, and/or *pCMV6-AN-Myc-Sobp*. 1.5 μg of each plasmid was transfected for all Co-IP assays. Proteins were extracted from HEK293T/17 cells after 48 hours using ice-cold Pierce IP Lysis Buffer (Thermo, 87787) with supplemental HALT Protease Inhibitor Cocktail with EDTA (Thermo, 1861281). Immunoprecipitation with either Pierce anti-C-Myc magnetic beads (Thermo, 88842), Pierce anti-HA magnetic beads (Thermo, 88836), or Pierce anti-DYKDDDDK magnetic beads (Thermo, A36797) was performed following the manufacturer’s protocol; with the exception that 1X tris buffered saline (TBS; Thermo, BP2471-1) with 1% Triton X-100 (TBST) was used as both the dilution and washing buffer for both the Pierce anti-HA and anti-DYKDDDDK magnetic beads. Bound proteins were washed five times, eluted using the Pierce Lane Marker Non-Reducing Sample Buffer (Thermo, 39001), and subsequently reduced using 100 mM DTT prior to SDS-PAGE and Western blot. For control experiments, each sample in IP Lysis Buffer was diluted with Laemmli Sample Buffer (Bio-Rad, 161-0747) and reduced with 2% β-mercaptoethanol (BME; Bio-Rad, 161-0710) prior to SDS-PAGE and Western blot.

Immobilon-FL PVDF membranes (Merck Millipore, IPFL00010) were probed with a combination of rabbit anti-SIX1; either mouse anti-Myc or rabbit anti-Myc to detect Myc-PA2G4, Myc-MCRS1, and Myc-SOBP; either mouse anti-HA or rabbit anti-HA to detect HA-EYA1 and HA-EYA2; and rabbit anti-β-actin (Cell Signaling, 13E5) as a control. All primary antibodies were diluted to a 1:1000 concentration. Secondary antibodies (IRDye 680RD donkey anti-rabbit IgG (Licor, 92568073) or IRDye 800CW goat anti-mouse IgG (Licor, 92532210)) were used at a concentration of 1:5,000 as appropriate. Experiments were repeated at least three independent times. Blots were scanned using the Licor Odyssey infrared imaging system. For each Co-IP assay, the reverse assay was also performed to validate the results were not a result of non-specific binding (data not shown).

### Luciferase assay

Transfected MC3T3-E1 cells were harvested and analyzed using the Dual Luciferase Assay kit (Promega, E1910) following the manufacturer’s protocol. Each transfection included 200 ng of *pGL3-6xMEF3-Firefly* luciferase reporter (Ford et al., 2000) and 20 ng of *Renilla* luciferase reporter (*pCMV-Renilla*), in addition to different combinations of *pCMV-AN-Myc* (control), *pCMV2-5’Flag-Six1*, *pHM6-HA-Eya1*, *pHM6-HA-Eya2*, *pCMV6-AN-Myc-Pa2G4*, *pCMV6-AN-Myc-Mcrs1*, and/or *pCMV6-AN-Myc-Sobp* (400 ng each). 48 hours post-transfection, cells were harvested in 1X Passive Lysis Buffer and stored at −80°C overnight to further facilitate cell lysis. 20 μL of lysate was used in the analysis. Experiments were repeated at least five independent times and read on a Varioskan Flash Plate Reader (Thermo). Paired ANOVA with Tukey’s post-hoc multiple comparisons test was performed using GraphPad Prism 9 software. Expression of exogenous proteins from the transfected plasmid was confirmed by standard Western blotting using the antibodies described for Co-IP.

### Quantitative real-time PCR (qPCR)

E10.5 mouse embryos were harvested in PBS and stored in RNAlater (Thermo; AM7020). After genotyping, the mandibular arch of control (*Six1^+/+^*) and *Six1^-/-^*were dissected, and three arches pooled for RNA extraction using the Direct-zol RNA Micro kit (Zymo; R2062). Untransfected MC3T3-E1 cells were cultured for 48 hours prior to RNA extraction using the Direct-zol RNA Micro kit. cDNA was synthesized using the iScript Advanced cDNA Synthesis kit (Bio-Rad, 1725037) and qPCR performed with 5 ng of cDNA and the Sso Advanced Universal SYBR Green Supermix (Bio-Rad, 1725270) in a Bio-Rad CFX-384 Real Time Thermal Cycler. QuantiTect Assay primers for *Actb* (housekeeping gene), *Six1*, *Eya1*, *Eya2*, *Pa2G4*, *Mcrs1*, and *Sobp* were purchased from Qiagen. All assays were performed in duplicates at least three times. Statistical analysis between control and *Six1^-/-^* embryos was performed with GraphPad Prism 9 and significance was calculated using an unpaired two-tailed t-test. Analysis of expression levels for *Six1* and co-factors in MC3T3-E1 cells was performed by plotting the threshold cycle (Ct) for each gene relative to the Ct for *Actb*.

## Results

### Pa2G4, Mcrs1, and Sobp are expressed in the mouse mandibular arch at E10.5 and *Six1* is required for the regulation of both *Pa2G4* and *Sobp*

As Six1 is critical for jaw development in mouse (Luo et al., 2023; Tavares et al., 2017; Guo et al., 2011) and Pa2G4, Mcrs1, and Sobp were identified as Six1 co-factors during otic and branchial cartilage development in *Xenopus laevis* (Neilson et al., 2017, Neilson et al., 2020, Tavares et al., 2021, Keer et al., 2022), it was important to establish whether these genes are co-expressed with *Six1* and its two well-established co-activators *Eya1* and *Eya2* (Ozaki et al., 2004, Tavares et al., 2017, Zhang et al., 2021) in the mandibular arch at embryonic day (E) 10.5 using whole mount *in situ* hybridization (ISH). E10.5 was selected because this is a timepoint by which the NCCs have migrated into the mandibular arch and are in the midst of specification towards to both cartilaginous and skeletal fates as a result of signals originating from the ectoderm and endoderm (Tucker et al., 1999, Clouthier et al., 2000, Alexander et al., 2011, Zuniga et al., 2011, Tavares et al., 2012, Seto et al., 2024). *Six1* (**Fig. 1A**) is expressed throughout the entire length of the oral half of the mandibular arch and is co-expressed with each of *Eya1* (**Fig. 1B**), *Eya2* (**Fig. 1C**), *Pa2G4* (**Fig. 1D**), *Mcrs1* (**Fig. 1E**), and *Sobp* (**Fig. 1F**). *Eya1* (**Fig. 1B**) is expressed primarily along the oral half of the mandibular arch, extending from the distal edge, where the two mandibular arches meet, a little over halfway to the proximal side of the mandibular arch.

**Figure 1:**
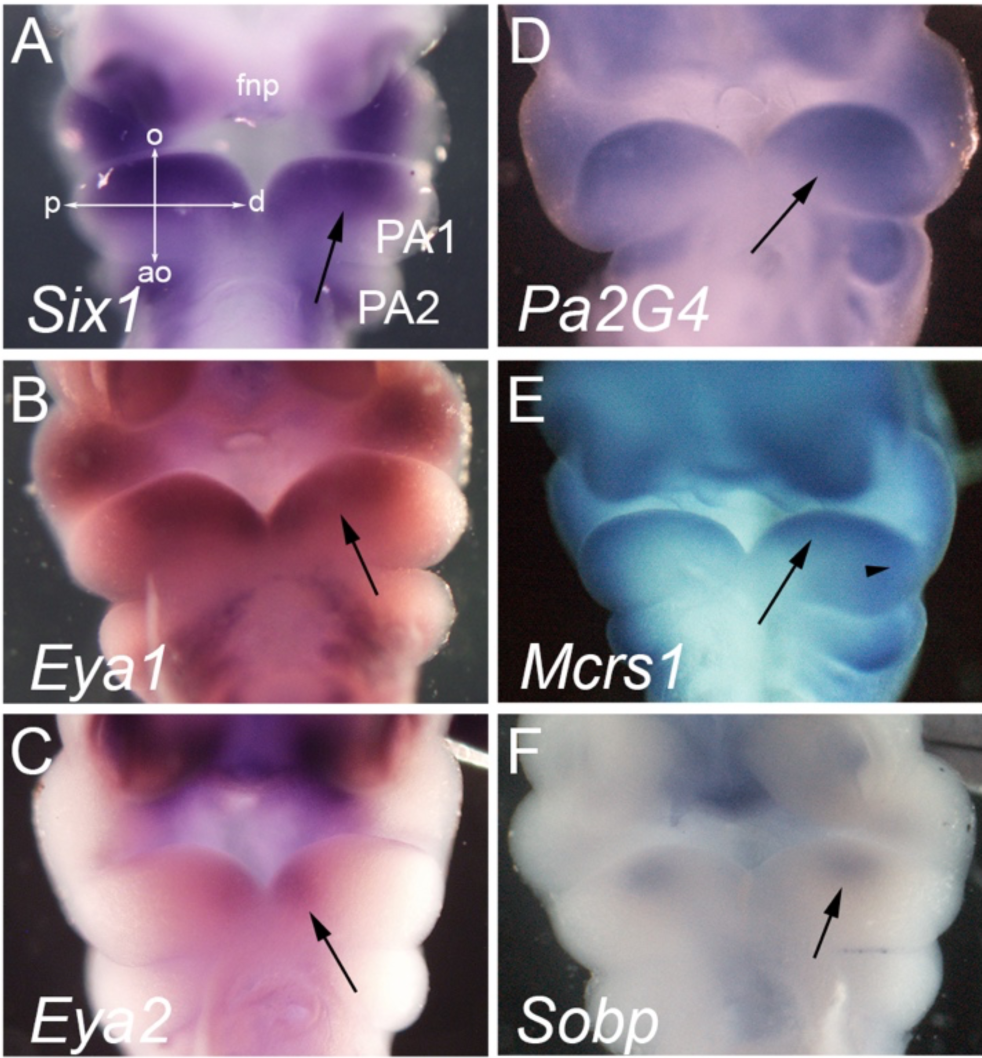
Expression domains of *Eya1*, *Eya2*, *Pa2G4*, *Mcrs1*, and *Sobp* overlap that of *Six1* in the first pharyngeal arch. Representative ventral view images of E10.5 mouse embryos after *in situ* hybridization for *Six1* (**A**), *Eya1* (**B**), *Eya2* (**C**), *Pa2G4* (**D**), *Mcrs1* (**E**), and *Sobp* (**F**). The heart was removed to better visualize the pharyngeal arches. **A**) *Six1* is expressed throughout the oral half of the mandibular arch (arrow), extending from the distal domain (where the two mandibular arches meet) through to the proximal domain. **B**) *Eya1* is expressed throughout the oral half of the mandibular arch, extending from the distal domain (arrow) approximately halfway across the arch and then tapering in expression toward the proximal domain, where there is little to no expression of *Eya1*. **C**) *Eya2* is expressed only in the distal domain of the mandibular arch, contacting both the oral and aboral halves of the arch (arrow). **D**) *Pa2G4* is expressed throughout the entirety of the mandibular arch (arrow). **E**) *Mcrs1* is expressed most strongly in the oral half of the mandibular arch, extending the entire length of the arch. In the proximal region of the arch, the expression of *Mcrs1* begins to extend from the oral to the aboral domains, although this expression is not as strong as that in the oral half of the arch (arrowhead). **F**) *Sobp* is expressed in a very restricted region of the mandibular arch, only in the oral half of the arch approximately halfway between the proximal and distal domains, although it appears to be skewed slightly to the distal domain (arrow). **ao:** aboral, **d**: distal, **fnp**: frontonasal prominence, **o**: oral, **p**: proximal, **PA1**: first pharyngeal arch (*i.e.*, mandibular arch), **PA2**: second pharyngeal arch.

*Eya2*, on the other hand, is primarily expressed along the distal edge of the mandibular arch, extending from the oral to the aboral half of the arch (**Fig. 1C**). Of the *Xenopus*-identified *Six1* co-factors, *Pa2G4* (**Fig. 1D**) is the most broadly expressed, covering almost the entire mandibular arch. *Mcrs1* (**Fig. 1E**) is largely expressed along the oral half of the mandibular arch, spanning the entire length of the arch and towards the proximal side of the arch, extends into the aboral region as well. *Sobp* (**Fig. 1F**) has the most restricted expression domain, located solely along the oral half of the mandibular arch, approximately halfway between the proximal and distal domains of the arch, although it appears to be slightly skewed towards the distal domain.

We also performed qPCR on mRNA isolated from the mandibular arch at E10.5 to determine whether the co-expression of *Six1* and putative co-factors *Pa2G4*, *Mcrs1*, and *Sobp* were biologically relevant in the developing arch. Analysis of the qPCR data reveals that, in the absence of *Six1* in *Six1^-/-^* mice, there was a significant reduction in the mRNA of *Pa2G4* and *Sobp* but not *Mcrs1*, suggesting that SIX1 is required for the expression of both *Pa2G4* and *Sobp* (**Fig. 2**).

**Figure 2:**
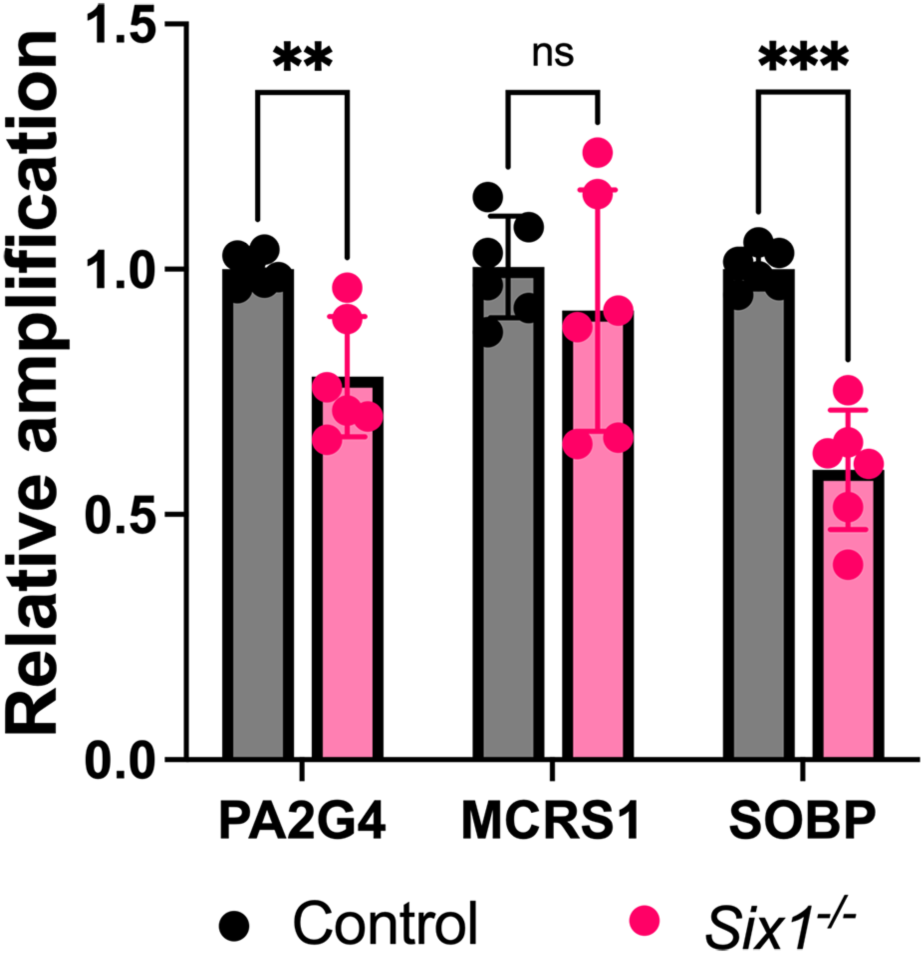
SIX1 is required for expression of *Pa2G4* and *Sobp* in the first pharyngeal arch. qPCR analysis of dissected mandibular arches from E10.5 control (*Six1^+/+^*) and *Six1^-/-^* mouse embryos demonstrate a significant decrease in mRNA for *Pa2G4* (*p* = 0.006090) and *Sobp* (*p* = 0.000208) after complete loss of SIX1 (*Six1^-/-^*). *Mcrs1* mRNA levels do not show a significant change (*p* = 0.438724). Data is shown relative to control embryos and normalized using β-actin (*Actb*). **ns**: not significant. **\*\***: *p* < 0.01; **\*\*\***: *p* < 0.001.

As whole mount ISH only indicates where mRNA is expressed and does not provide any information as to the distribution of these proteins within the mandibular arch, sectional immunofluorescence (IF) was performed at E10.5 to determine which cell types co-express SIX1 and another co-factor. SIX1 is expressed in the ectomesenchyme throughout the oral half of the mandibular arch and is absent from the ectodermal epithelium in the distal (**Fig. 3B, D, L, N, V, X**), oral (**Fig. 3E, G, O, Q, Y, AA**) and proximal (**Fig. 3H, J, R, T, BB, DD**) domains. In agreement with the whole-mount ISH, PA2G4 has a broad expression pattern (**Fig. 3A**), as it is expressed throughout both the surface ectoderm and the ectomesenchyme in the distal (**Fig. 3C, D**) and oral (**Fig. 3F, G**) domains. In the proximal domain, while the ectomesenchyme continues to strongly express PA2G4, the fluorescence intensity in the proximal epithelium is reduced as compared to that of the oral and distal domains (**Fig. 3I, J**). MCRS1 appears to have a broad, low level expression pattern throughout both the ectoderm and ectomesenchyme, with small foci of increased expression within the ectomesenchyme (**Fig. 3K**). In the distal domain, MCRS1 is expressed primarily in the ectoderm and has a broad, low-level expression throughout the ectomesenchyme (**Fig. 3M, N**) and strong expression in a few, scattered ectomesenchymal cells (**Fig. 3M**, arrowheads). In the oral domain, MCRS1 is primarily expressed in the ectomesenchymal cells subjacent to the ectoderm (**Fig. 3P, Q**). We also observed a small number of cells with slightly stronger expression of MCRS1 in the ectomesenchyme, most of which also strongly expressed SIX1 (**Fig. 3P**, arrowheads). In the proximal domain of the arch, MCRS1 expression appears to be more cytoplasmic and ubiquitous throughout both the ectoderm and the ectomesenchyme (**Fig. 3S, T**). As noted in the whole mount ISH (**Fig. 1F**), SOBP expression is limited within the mandibular arch (**Fig. 3U**). There was no obvious expression of SOBP within the distal domain of the arch (**Fig. 3W**, **X**). In the oral domain, we observed expression of SOBP within the ectoderm (**Fig. 3Z**, arrow) and in the ectomesenchyme subjacent to the ectoderm (**Fig. 3Z**, arrowheads). It is interesting to note that in the ectomesenchyme making direct contact with the ectoderm, there appears to be punctate expression of SOBP throughout the cytoplasm and the nuclei (**Fig. 3AA**). In the proximal domain of the arch, there was only strong expression of SOBP within the ectoderm (**Fig. 3CC**, arrow). Altogether, our ISH and IF findings demonstrate that *Pa2G4*, *Mcrs1*, and *Sobp* overlap with that of *Six1* and *Eya1*, primarily in the oral ectomesenchyme of the mandibular arch. Therefore, they are optimally located to regulate SIX1 function during NCC proliferation and differentiation within the mandibular arch and could contribute to the NCC-derivatives affected in patients with BOR. Of note, these proteins are also expressed in SIX1-negative areas of the mandibular arch, such as the surface ectoderm and aboral ectomesenchyme indicating that they may also have SIX1-independent roles.

**Figure 3:**
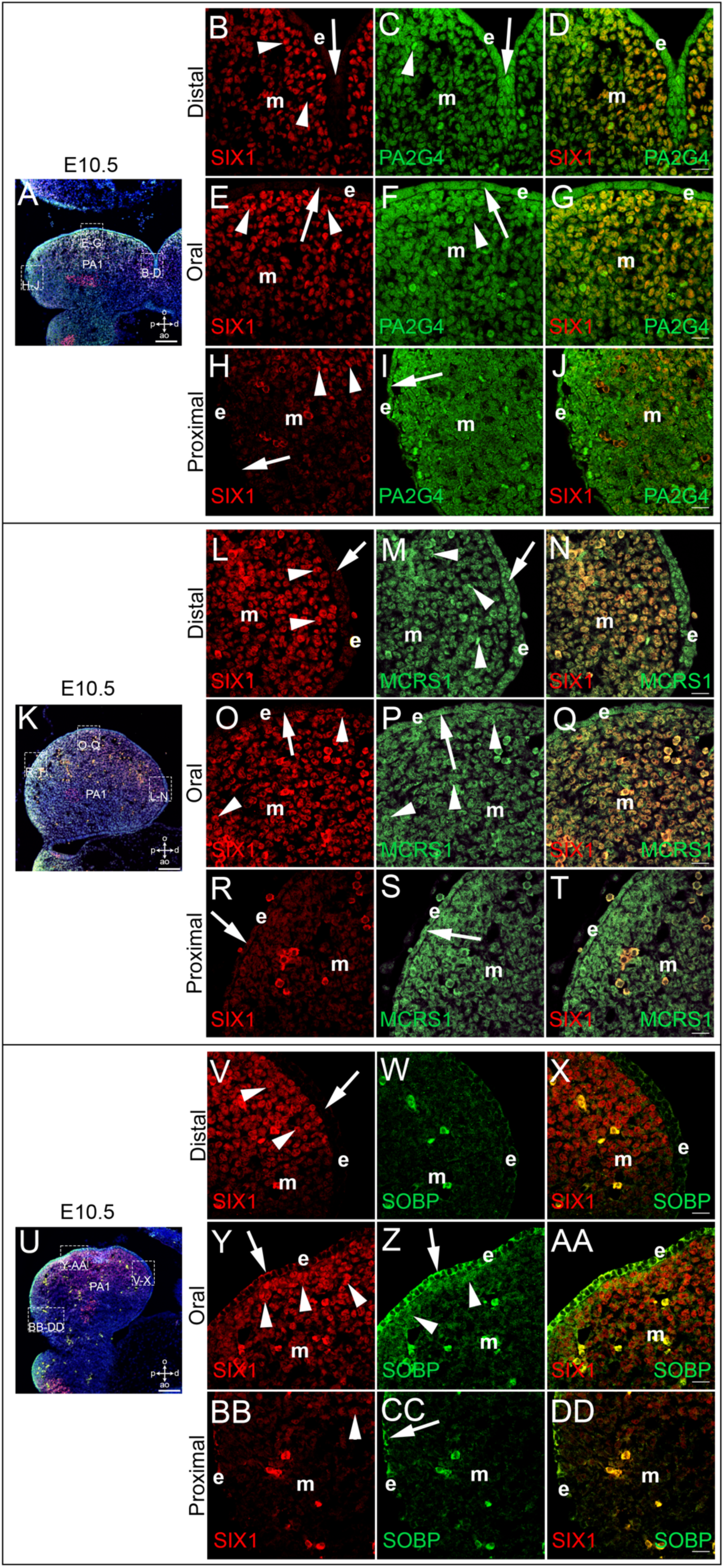
SIX1 is co-expressed with PA2G4, MCRS1, and SOBP within the mandibular arch. **A**, **K**, **U**) Representative lower magnification image of an E10.5 mandibular arch stained for SIX1 (red) and each respective co-factor (*i.e.*, PA2G4 (**A**), MCRS1 (**K**), and SOBP (**U**); green) showing the location of high magnification insets for the distal (**B**-**D**, **L-N**, **V-X**), oral (**E**-**G**, **O-P**, **Y-AA**), and proximal (**H**-**I**, **R-T**, **BB-DD**) domains of the arch. **B**, **E**, **H**, **L**, **O**, **R**, **V**, **Y**, **BB**) SIX1 is not expressed in the ectoderm in any of the samples examined (arrows). In the distal domain (**B**, **L**, **V**), SIX1 is strongly expressed in the ectomesenchyme subjacent to the oral ectoderm (arrowheads). In the oral domain (**E**, **O**, **Y**), SIX1 is strongly expressed in the ectomesenchyme subjacent to the ectoderm and then tapers off towards the aboral domain (arrowheads). In the proximal domain (**H**, **R**, **BB**), the expression of SIX1 has begun to taper off, with low-level expression in the ectomesenchyme. In the most oral of these proximal high-magnification insets (**R**), there is still some stronger expression of SIX1 in the ectomesenchyme (arrowheads) as compared to the more aboral high-magnification insets (**H**, **BB**), which have lower overall levels of SIX1 expression, with only medium expression in the most oral parts of the sections (arrowheads). PA2G4 (**C**, **D**, **F**, **G**, **I**, **J**) is broadly expressed throughout the mandibular arch and is expressed in both the epithelium (arrows) and the ectomesenchyme (arrowheads). In the distal domain (**C**), PA2G4 is most strongly expressed in the ectoderm and the ectomesenchyme subjacent to the ectoderm. In the oral domain (**F**), PA2G4 continues to be most strongly expressed in the ectoderm and the ectomesenchyme directly subjacent to the ectoderm, with scattered cells with strong expression of PA2G4. In the proximal domain (**I**), the expression pattern of PA2G4 appears to change, being expressed more in the cell membrane than the cytoplasm, though the expression is still strongest in the ectomesenchyme directly subjacent to the ectoderm, which has a lower overall expression of PA2G4 than the ectoderm in the other regions of the mandibular arch. Overall, PA2G4 is strongly co-expressed with SIX1 in the ectomesenchyme subjacent to the ectoderm in the oral half of the mandibular arch (**D**, **G**, **J**). MCRS1 (**M**, **N**, **P**, **Q**, **S**, **T**) is generally expressed throughout the mandibular arch at a low level in both the ectoderm (arrows) and the ectomesenchyme, although there are also scattered ectomesenchymal cells with stronger expression throughout the ectomesenchyme (arrowheads) in the distal (**M**) and oral (**P**) domains. The expression is largely the same throughout the proximal domain (**S**). Overall, MCRS1 is strongly co-expressed with SIX1 in the ectomesenchyme subjacent to the ectoderm in the oral half of the mandibular arch (**N**, **Q**, **T**). In agreement with the *in situ* hybridization data, SOBP (**W**, **X**, **Z**, **AA**, **CC**, **DD**) is only expressed in a very small region of the mandibular arch. In the distal domain (**W**), SOBP is only found in red blood cells and is absent from the epithelium and the ectomesenchyme. In the oral domain (**Z**), SOBP is strongly expressed in the ectoderm (arrow) and is also found in the ectomesenchyme directly subjacent to the ectoderm (arrowheads). Interestingly, in the proximal domain (**CC**), there was some expression in the ectoderm (arrow), but there was no notable expression in the ectomesenchyme. Overall, SOBP is strongly co-expressed with SIX1 in the ectomesenchyme of the oral domain of the arch (**AA**). **m**: ectomesenchyme, **e**: ectoderm, **ao**: aboral, **d**: distal, **o**: oral, **p**: proximal, **PA1**: first pharyngeal arch (*i.e.*, mandibular arch). DAPI was used as a nuclear counterstain (blue in **A**, **K**, and **U**). Scale bars are 100 μm in **A**, **K**, **U**; 20 μm in **D**, **G**, **J**, **N**, **Q**, **T**, **X**, **AA**, **DD**.

### SIX1 and MCRS1 are *bona fide* SIX1 co-factors in mouse

As the expression domains of the PA2G4, MCRS1, and SOBP proteins partially overlap with that of SIX1 in the mandibular arch, it is important to determine whether they are *bona fide* SIX1 co-factors in the mouse (*i.e.*, proteins that bind to SIX1 and modulate its transcriptional activity) as shown in *Xenopus* (Jourdeuil et al., 2023, Keer et al., 2022, Neilson et al., 2017, Neilson et al., 2020, Tavares et al., 2021). To assay whether these mouse proteins can bind to mouse SIX1, HEK293T cells were transfected with different combinations of control vector (*pCMV6-AN-Myc*), Flag-tagged *Six1*, Myc-tagged *Pa2G4*, Myc-tagged *Mcrs1*, or Myc-tagged *Sobp* followed by co-immunoprecipitation (Co-IP) assays. Western blot detection reveals that mouse PA2G4 does not bind to mouse SIX1 (**Fig. 4A**), whereas both MCRS1 (**Fig. 4B**) and SOBP (**Fig. 4C**) do.

**Figure 4:**
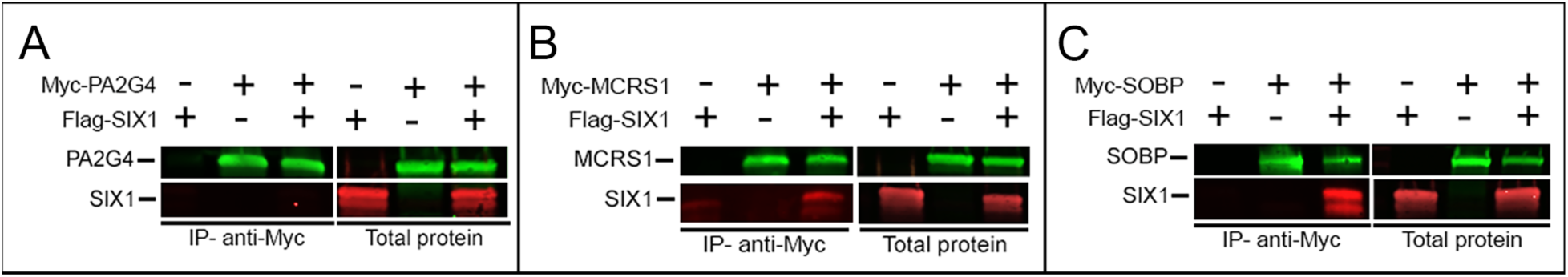
SIX1 binds to MCRS1 and SOBP. **A-C**) Multiplex fluorescent Western blot detection of SIX1 (red in **A**-**C**) and either Myc-PA2G4 (green in **A**), Myc-MRCS1 (green in **B**) or Myc-SOBP (green in **C**). In each panel (**A**-**C**), the data are organized such that the Western blot of the immunoprecipitation experiment (IP; using anti-c-Myc magnetic beads) is on the left and the Western blot of the total protein that was used in the IP experiment is on the right. Each Western blot is organized such that the first lane (Myc -, Six1 +) represents the protein generated from cells transfected only with Flag-tagged SIX1, the second lane (Myc +, Six1 -) represents the protein generated from cells transfected only with a Myc-tagged co-factor (*i.e.*, PA2G4, MCRS1, or SOBP), and the third lane (Myc +, Six1 +) represents the protein generated from cells transfected with both a Myc-tagged co-factor and Flag-tagged SIX1. The results show both that the appropriate proteins were present in each sample prior to the IP experiment and that after IP with c-Myc magnetic beads PA2G4 (**A**) is unable to bind to or precipitate SIX1 while both MCRS1 (**B**) and SOBP (**C**) do bind and precipitate SIX1. The inverse experiments, in which IP was performed with Flag-tagged magnetic beads were also performed as a control and generated the same results (not shown).

To determine whether these proteins are then able to modulate the transcriptional activity of SIX1, standard luciferase assays using MC3T3-E1 cells were performed. These analyses reveal that in these cells, SIX1 alone drives luciferase activity at a significantly higher level than that of the control vector (**Fig. 5A**). This is likely because MC3T3-E1 cells express endogenous EYA1 (**Supplemental Fig. 2**), which can activate SIX1 transcriptional activity. However, when PA2G4, MCRS1, or SOBP are co-transfected with SIX1, there is no significant different in the luciferase activity when compared to that of SIX1 alone (**Fig. 5A**). Consistent with previous reports (Li et al., 2003, Shah et al., 2020, Tavares et al., 2021, Patrick et al., 2009), co-transfection of SIX1+EYA1 results in an increase in luciferase activity (**Fig. 5B**). Surprisingly, when PA2G4, MCRS1, and SOBP were co-transfected with SIX1+EYA1 to determine whether they were able to modulate the transcriptional activity of the SIX1+EYA1 complex as has been described in *Xenopus* (Keer et al., 2022, Neilson et al., 2017, Neilson et al., 2020, Tavares et al., 2021), neither PA2G4 nor MCRS1 significantly affect the activity of the SIX1+EYA1 transcriptional complex, whereas SOBP significantly represses it (**Fig. 5B**). However, as EYA2 can also act as a SIX1 co-activator (Patrick et al., 2009), is expressed in the mandibular arch (**Fig. 1C**), and is expressed at low levels in the MC3T3-E1 cells (**Supplemental Fig. 2**), we tested whether these three proteins are also able to modulate the transcriptional activity of the SIX1+EYA2 complex. Interestingly, we detected a significantly larger increase in the luciferase activity of the reporter vector after co-transfection of *Six1+Eya2* compared to that of *Six1+Eya1* (*p* < 0.0001). PA2G4 significantly enhances SIX1+EYA2 transcriptional activity, whereas both MCRS1 and SOBP significantly represses it (**Fig. 5C**). Altogether, these results demonstrate that MCRS1 and SOBP are *bona fide* SIX1 co-repressors. Interestingly, in the mouse, PA2G4 acts as an enhancer of the SIX1+EYA2 transcriptional complex without binding to SIX1.

**Figure 5:**
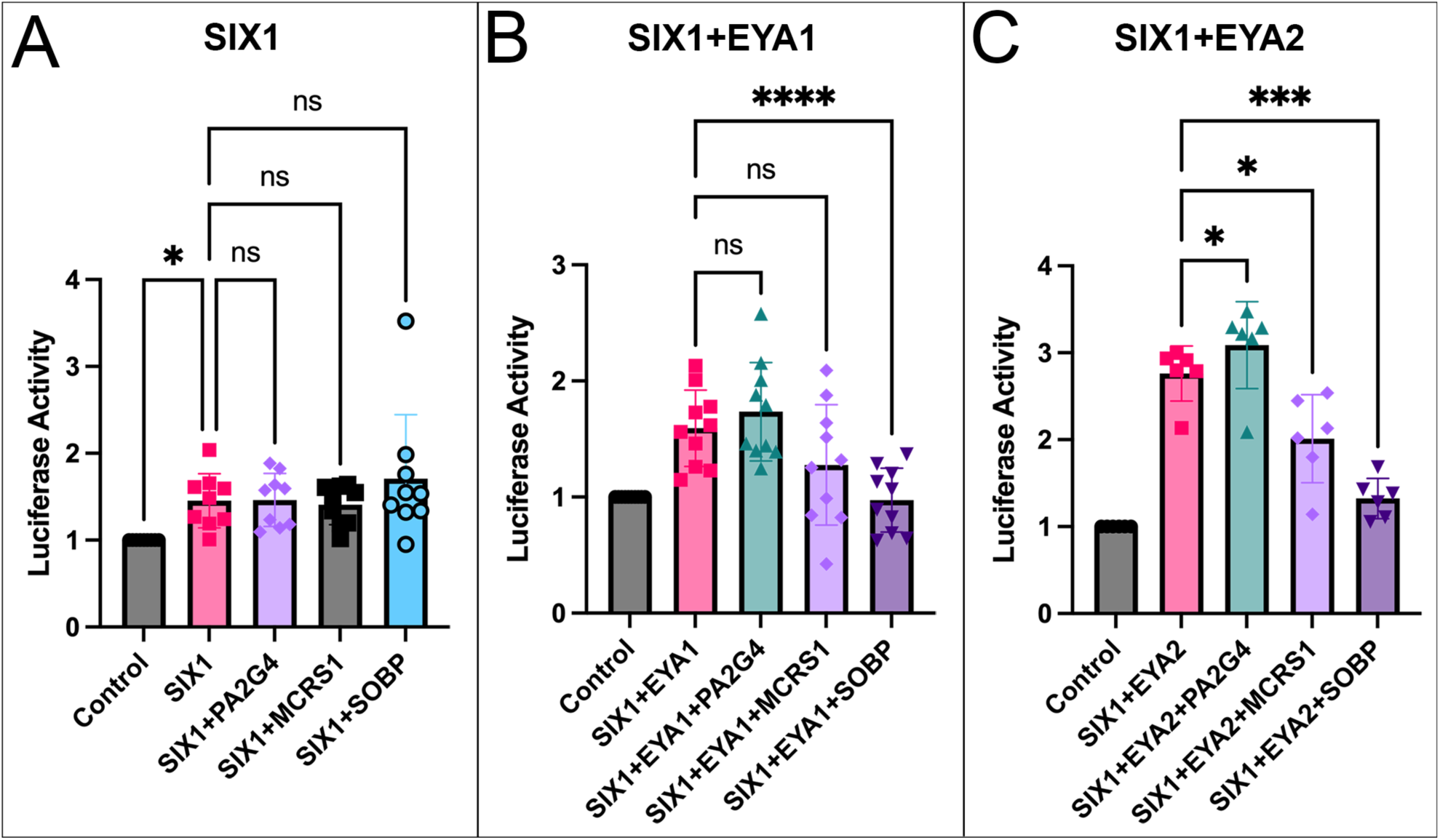
MCRS1 and SOBP repress SIX1+EYA transcriptional activity while PA2G4 enhances it. **A**-**C**) Graphs depicting the luciferase activity of the pGL3-6xMEF3-luciferase reporter in MC3T3-E1 cells transfected with different combinations of either control vector, *Flag-Six1*, *Ha-Eya1*, *HA-Eya2*, *Myc-Pa2G4*, *Myc-Mcrs1*, and/or *Myc-Sobp*. Data were normalized to CMV-Renilla and are represented relative to control cells. **A**) Luciferase activity of the reporter vector is significantly increased after expression of SIX1 relative to control (*p* < 0.05) whereas there is no significant difference in luciferase activity after expression of *Pa2G4*, *Mcrs1*, or *Sobp* relative to *Six1* (ns). **B**) After expression of *Six1*+*Eya1*, there is a significant increase in the luciferase activity of the reporter vector (*p* < 0.001). Expression of *Six1*+*Eya1* with either *Pa2G4* or *Mcrs1* does not significantly affect the activity of the reporter vector relative to *Six1*+*Eya1* (ns). Expression of *Sobp* with *Six1*+*Eya1*, on the other hand, significantly (*p* < 0.0001) represses the luciferase activity of the reporter vector relative to *Six1*+*Eya1*. **C**) There is a significantly larger increase in luciferase activity of the reporter vector after expression of *Six1*+*Eya2* compared to that of *Six1*+*Eya1* (*p* < 0.0001). Expression of *Pa2G4* with *Six1*+*Eya2* significantly enhances the activity of the reporter vector relative to *Six1*+*Eya2* (*p* < 0.05) while expression of *Mcrs1* (*p* < 0.05) or *Sobp* (*p* < 0.001) with *Six1*+*Eya2* significantly represses the luciferase activity of the reporter vector relative to *Six1*+*Eya2*.

### SIX1 translocates MCRS1 to the nucleus

In order to bind to SIX1, the co-activator EYA, which is a cytoplasmic protein, must be translocated either by SIX1 or other co-factors to the nucleus (Li et al., 2003, Ohto et al., 1999). Therefore, we sought to determine whether the subcellular localization of PA2G4, MCRS1, or SOBP change when co-transfected with SIX1 in MC3T3-E1 cells. First, as expected from previous reports (Ohto et al., 1999, Tavares et al., 2021, Shah et al., 2020), when transfected singly SIX1 is localized to the nucleus (**Fig. 6A-C**) whereas EYA1 is cytosolic (**Fig. 6D-F**).

**Figure 6:**
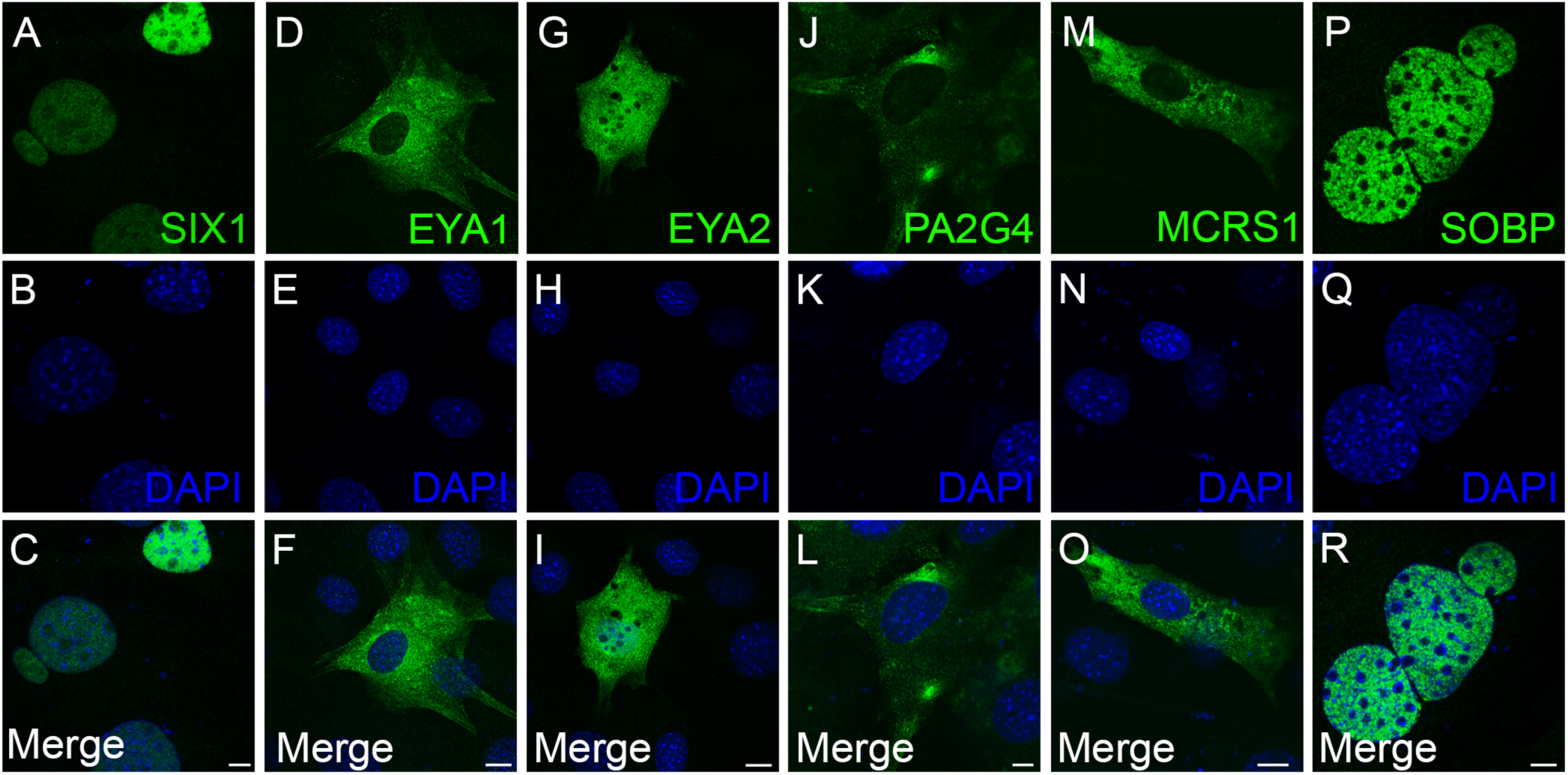
Subcellular localization of SIX1, EYA1, EYA2, PA2G4, MCRS1, and SOBP. Confocal images of MC3T3-E1 cells expressing *Flag-Six1* (**A**, **C**), *HA-Eya1* (**D**, **F**), *HA-Eya2* (**G**, **I**), *Myc-Pa2G4* (**J**, **L**), *Myc-Mcrs1* (**M**, **O**), or *Myc-Sobp* (**P**, **R**) are shown in green while the nucleus (**B**-**C, E**-**F**, **H**-**I**, **K**-**L**, **N**-**O**, **Q**-**R**), which are counterstained with DAPI, are shown in blue. **A**-**C**) SIX1 is expressed in the nucleus. **D**-**F**) Eya1 is expressed in the cytoplasm. **G**-**I**) EYA2 is primarily expressed throughout the cytoplasm, with some nuclear expression. **J**-**L**) PA2G4 is expressed in the cytoplasm and nucleolus. **M**-**O**) MCRS1 is expressed in the cytoplasm. **P**-**R**) SOBP is expressed in the nucleus. Scale bars: 10 μm in **C**, **F**, **I**, **R**; 5 μm in **L**, **O**.

EYA2 was located throughout both the cytosol and nucleus (**Fig. 6G-I**). PA2G4 was primarily cytosolic with some nucleolar staining (**Fig. 6J-L**) while MCRS1 was mainly cytosolic (**Fig. 6M-O**). SOBP, as previously reported (Tavares et al., 2021), is solely nuclear (**Fig. 6P-R**). When SIX1 and EYA1 are co-transfected, EYA1 is fully translocated to the nucleus (**Fig. 7A-D**), which agrees with previous reports (Ohto et al., 1999, Tavares et al., 2021, Shah et al., 2020). We also observed a complete translocation of EYA2 to the nucleus when co-transfected with SIX1 (**Fig. 7E-H**). Co-transfection of SIX1 and PA2G4 did not alter the subcellular localization of either SIX1 or PA2G4 (**Fig. 7I-L**). In contrast, co-expression of SIX1 and MCRS1 resulted in a partial translocation of MCRS1 from the cytoplasm to the nucleus (**Fig. 7M-P**). When SIX1 and SOBP are co-transfected, there was no change in the subcellular localization of either protein, as both were already nuclear (**Fig. 7Q-T**). These data further corroborate that MCRS1 is a *bona fide* SIX1 co-factor in the mouse, as it can both bind to and be translocated to the nucleus by SIX1, in agreement with what has been reported in *Xenopus* for Sobp (Tavares et al., 2021).

**Figure 7:**
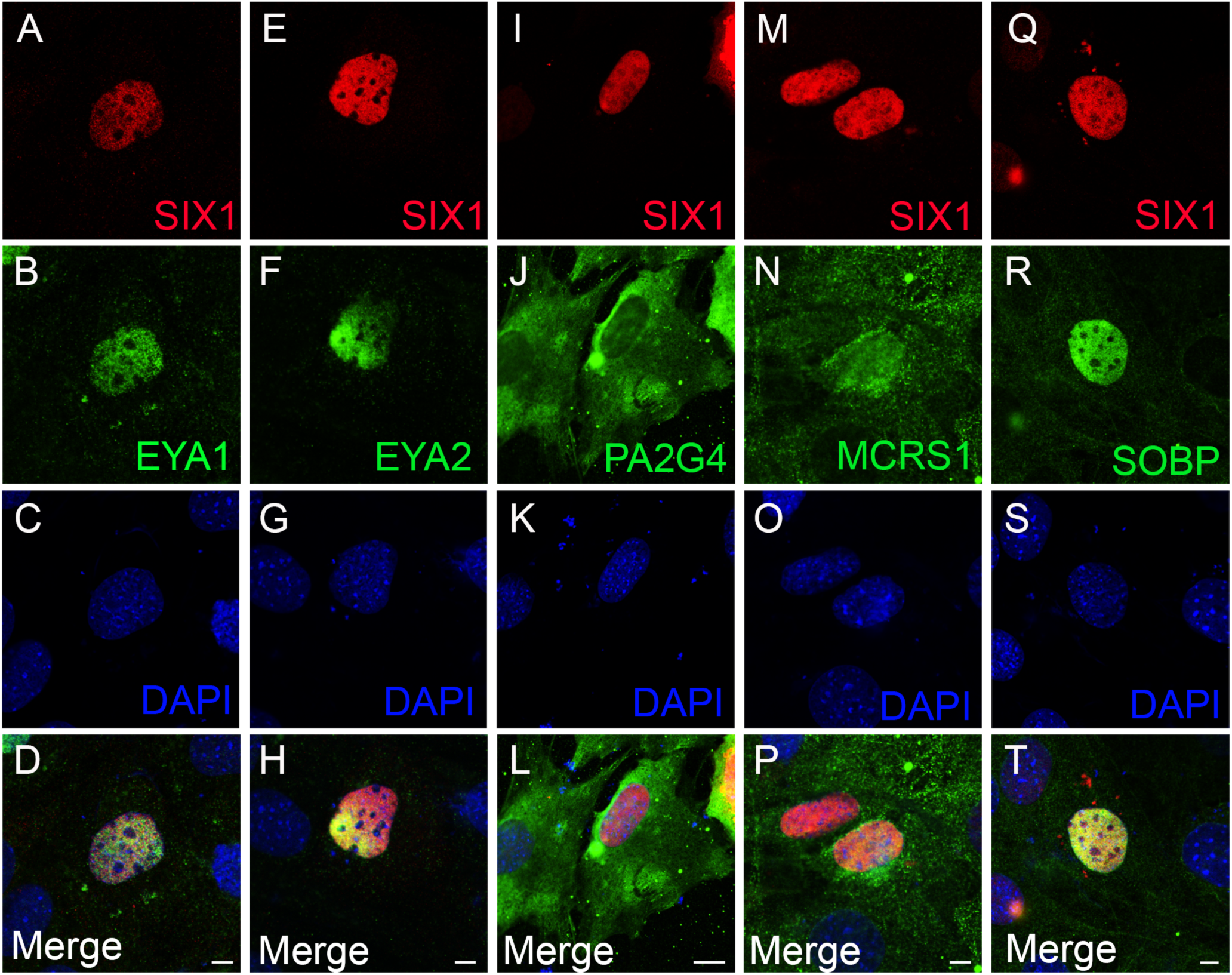
SIX1 translocates EYA1, EYA2, and MCRS1 to the nucleus. Confocal images of MC3T3-E1 cells co-expressing *Flag-Six1* (red in **A**, **D**, **E**, **H**, **I**, **L**, **M**, **P**, **Q**, **T**) and one of *HA-Eya1* (**B**, **D**), *HA-Eya2* (**F**, **H**), *Myc-Pa2G4* (**J**, **L**), *Myc-Mcrs1* (**N**, **P**), or *Myc-Sobp* (**R**, **T**) in green. For each cell the nucleus (**C-D**, **G-H**, **K-L**, **O-P**, **S-T**) was identified with DAPI and shown in blue. **A**-**D**) SIX1 translocates EYA1 from the cytoplasm to the nucleus. **E**-**H**) SIX1 translocates EYA2 from the cytoplasm to the nucleus. **I**-**L**) There is no change in the subcellular localization of either PA2G4 (cytoplasm and nucleolus) or SIX1 (nucleus). **M**-**P**) Six1 translocates some, but not all, of the cytoplasmic MCRS1 to the nucleus. **Q**-**T**) There is no change in the subcellular localization of either SIX1 (nucleus) or SOBP (nucleus). Scale bars: 5 μm in **D**, **H**, **P**, **T**; 10 μm in **L**.

### MCRS1 and PA2G4 indirectly translocate EYA to the nucleus

Tavares *et al*. (2021) demonstrated that, in *Xenopus*, Sobp represses the transcriptional activity of the Six1+Eya1 complex by competing with Eya1 for binding to Six1 and, in turn, translocating Eya1 from the nucleus to the cytosol. Further, we have shown that, in mouse, PA2G4 is able to enhance the transcriptional activity of the SIX1+EYA complex without directly binding to SIX1 (**Fig. 5C**). Therefore, we hypothesized that PA2G4, MCRS1, and SOBP may be able to indirectly regulate the activity of the SIX1+EYA transcription complex by translocating EYA to the nucleus and making it available to SIX1. To test this, either EYA1 or EYA2 were co-transfected with PA2G4, MCRS1, or SOBP in MC3T3-E1 cells to determine whether their co-expression altered the subcellular localization of these proteins as described above. When PA2G4 and EYA1 were co-transfected (**Fig. 8A-D**), we observed a partial translocation of EYA1 to the nucleus while PA2G4 remains largely cytosolic. Co-transfection of MCRS1 and EYA1 (**Fig. 8E-H**) results in a partial translocation of EYA1 and MCRS1 to the nucleus. Interestingly, co-transfection of EYA1 and SOBP resulted in a partial translocation of EYA1 to the nucleus and a partial translocation of SOBP to the cytosol (**Fig. 8I-L**). In addition, both proteins appeared to be more punctate in expression, as compared to their normally “smooth” appearance. This was not an artifact of the staining as the punctate expression appeared in all co-transfected cells on all slides examined and these proteins did not have this punctate expression in any other conditions examined during this study. Co-transfection experiments with EYA2 yielded slight differences in the subcellular distribution of all proteins as compared to those with EYA1 (compare **Fig. 8A-L** to **8M-X**). Co-transfection of EYA2 and PA2G4 resulted in a partial translocation of EYA2 to the nucleus, however this was less effective than when EYA1 and PA2G4 are co-expressed, as more EYA2 remains in the cytosol (**Fig. 8M-P**). Co-transfection of EYA2 and MCRS1 results in an almost complete translocation of both EYA2 and MCRS1 to the nucleus, with some faint expression of each remaining in the cytosol (**Fig. 8Q-T**). Co-transfection of SOBP and EYA2 results in a complete translocation of EYA2 to the nucleus and no coincident translocation of SOBP to the cytosol (**Fig. 8U-X**). These results further corroborate the mechanism previously reported in *Xenopus* for Sobp (Tavares et al., 2021) whereby modulation of SIX1 transcriptional activity by some Six1 co-factors may be achieved by altering the subcellular localization of Eya proteins. Further, these data suggest that the interactions between PA2G4, MCRS1, and SOBP are different based on the EYA family member with which they are interacting.

**Figure 8:**
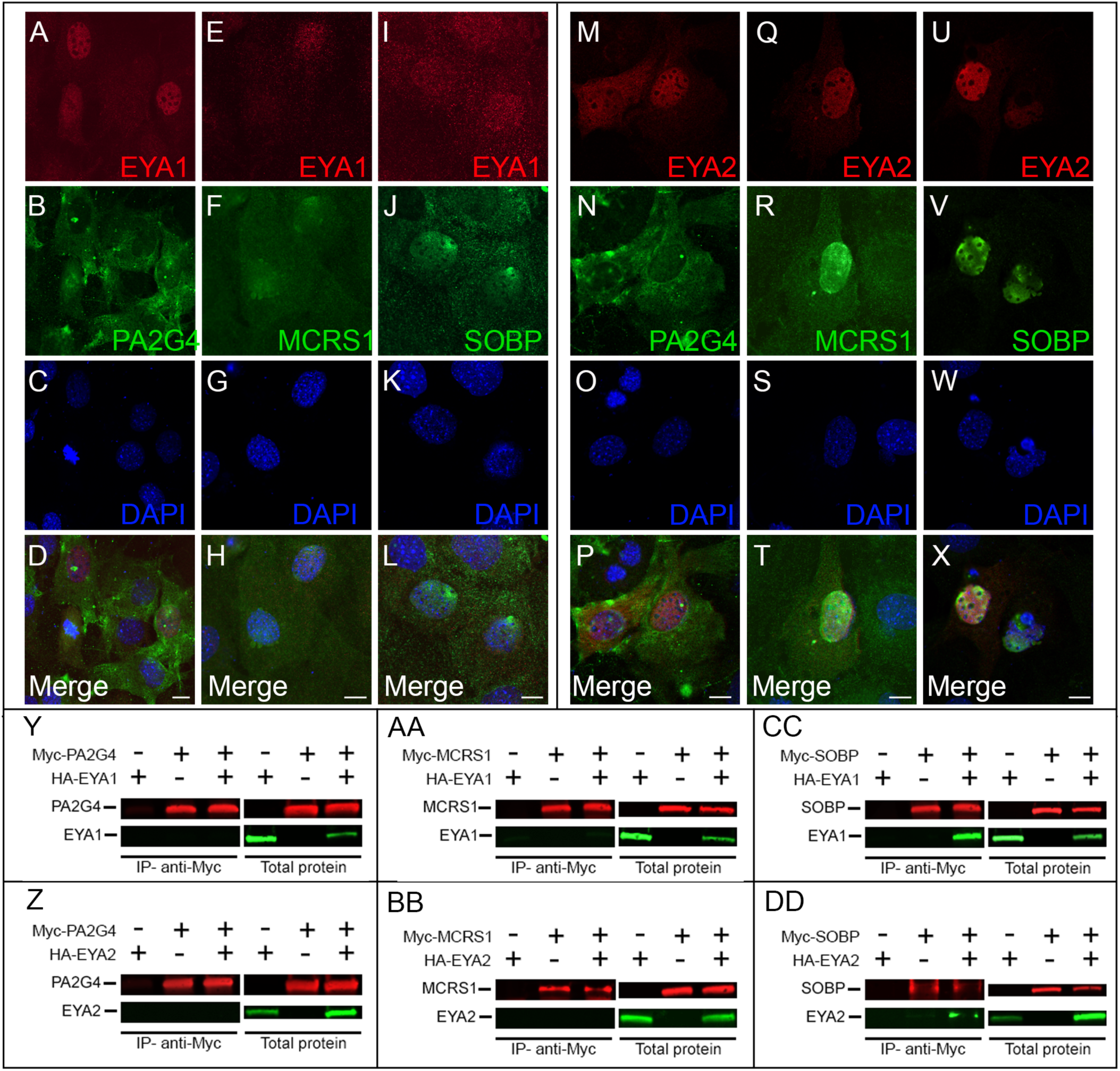
PA2G4, MCRS1, and SOBP alter the subcellular localization of EYA1 and EYA2 via indirect (PA2G4 and MCRS1) or direct (SOBP) binding to EYA proteins. **A**-**X**) Confocal images of MC3T3-E1 cells co-transfected with either *HA-Eya1* (**A**, **D-E**, **H-I**, **L**) or *HA-Eya2* (**M**, **P-Q**, **T-U**, **X**) shown here in red and one of either *Myc-Pa2G4* (**B**, **D**, **N**, **P**), *Myc-Mcrs1* (**F**, **H**, **R**, **T**), or *Myc-Sobp* (**I**, **L**, **V**, **X**) shown here in green. For each cell, the nucleus (**C**-**D**, **G**-**H**, **K**-**L**, **O**-**P**, **S**-**T**, **W**-**X**) was also identified with DAPI, shown here in blue. **A**-**D**) When PA2G4 is co-expressed with EYA1, the subcellular localization of PA2G4 is unchanged while EYA1 is partially translocated to the nucleus. **E**-**H**) When MCRS1 and EYA1 are co-expressed, both MCRS1 and EYA1 are partially translocated to the nucleus. **I**-**L**) When SOBP is co-expressed with EYA1, the expression of both SOBP and EYA1 become punctate and curiously SOBP is partially translocated to the cytoplasm while EYA1 is partially translocated to the nucleus. **M**-**P**) When PA2G4 and EYA2 are co-expressed, EYA2 is partially translocated to the nucleus while the expression of PA2G4 remained unchanged and is found only in the cytoplasm. **Q**-**T**) When MCRS1 is co-expressed with EYA2, EYA2 is partially translocated to the nucleus, while MCRS1 was largely translocated to the nucleus although some cytoplasmic expression remains. **U**-**X**) When SOBP and EYA2 are co-expressed, EYA2 is almost entirely translocated to the nucleus while SOBP remained exclusively nuclear. Scale bars: 10 μm. In (**Y**-**DD**) Multiplex fluorescent Western blot detection of either HA-EYA1 (green in **Y, AA**, **CC**) or HA-EYA2 (green in **Z**, **BB**, **DD**) with one of MYC-PA2G4 (red in **Y**, **Z**), MYC-MCRS1 (red in **AA**, **BB**), or MYC-SOBP (red in **CC**, **DD**). In each panel (**Y**-**DD**), the data are organized such that the Western blot of the immunoprecipitation experiment (IP; using anti-c-Myc magnetic beads) is on the left and the Western blot of the total protein that was used in the IP experiment is on the right. Each Western blot is organized such that the first lane (Myc -, HA +) represents the protein generated from cells transfected only with HA-tagged EYA1 or EYA2, the second lane (Myc +, HA -) represents the protein generated from cells transfected only with a Myc-tagged co-factor (*i.e.*, PA2G4, MCRS1, or SOBP), and the third lane (Myc +, HA +) represents the protein generated from cells transfected with both a Myc-tagged co-factor and an HA-tagged EYA. The results show both that the appropriate proteins were present in each sample prior to the IP experiment and that after IP with c-Myc magnetic beads PA2G4 is unable to bind or precipitate either EYA1 (**Y**) or EYA2 (**Z**). MCRS1 was also unable to either bind or precipitate EYA1 (**AA**) or EYA2 (**BB**). SOBP, on the other hand was able to bind and precipitate both EYA1 (**CC**) and EYA2 (**DD**). The inverse experiments, in which IP was performed with HA-tagged magnetic beads were also performed as a control and generated the same results (not shown).

Tavares *et al*. (2021) demonstrated that the nuclear to cytosolic translocation of Eya1 by Sobp in *Xenopus* occur via direct binding between Eya1 and Sobp without the addition of exogenous Six1. As we have observed that PA2G4, MCRS1, and SOBP have some ability to translocate EYA proteins to the nucleus and further, that PA2G4 enhances the transcriptional activity of the SIX1+EYA2 complex without binding directly to SIX1, it was important to determine whether PA2G4, MCRS1, or SOBP are able to bind to EYA1/2 in mouse. Co-IP experiments revealed that neither PA2G4 (**Fig. 8Y-Z**) nor MCRS1 (**Fig. 8AA-BB**) bind to EYA1 or EYA2. SOBP, on the other hand, binds to both EYA1 (**Fig 8CC**) and EYA2 (**Fig. 8DD**), in agreement with what has previously been reported in *Xenopus* for Eya1 (Tavares et al., 2021).

### SOBP and MCRS1 are co-expressed with SIX1 during craniofacial bone development

A recent study by Baxi et al. (2023) revealed that *Six1* regulates genes in the mandibular arch related to NCC patterning and bone development. This underscores the importance of identifying whether these *bona fide* SIX1 co-factors are co-expressed with SIX1 during later craniofacial bone development. To investigate this, we collected mouse embryos at E14.5 and performed immunohistochemistry for SIX1 and SOBP on transverse sections through the developing mandible and incisors. This is a period of ongoing differentiation of ectomesenchyme into both osteoblasts and, as these osteoblasts become buried in extracellular matrix, osteocytes (Franz-Odendaal et al., 2006). Transverse sections through the developing incisors (**Fig. 9A-C**) and mandible (**Fig. 9D-F**) demonstrate that SIX1 is expressed within the oral epithelium, dental lamina, dental follicle and at the boundary of the condensed ectomesenchyme surrounding the developing incisor (**Fig. 9A, C**). Within the developing mandible, we noted increased expression of SIX1 within the periosteum of the developing mandible and within the condensed ectomesenchyme surrounding the developing incisor. We also noted some expression within Meckel’s cartilage (**Fig. 9D, F**). Interestingly, while SOBP was largely co-expressed with SIX1 in these tissues, there were distinct regions with expression of only SOBP or SIX1 (**Fig. 9C, F**). SOBP was strongly expressed within the dental follicle and had a weak and asymmetrical distribution within the dental lamina and the stellate reticulum (**Fig. 9B**). There was then an area of low to no SOBP expression within the condensed ectomesenchyme surrounding the developing incisor and then strong co-expression of SOBP and SIX1 within the boundary of the condensed mesenchyme (**Fig. 9B, C**). Importantly, the area of strong SOBP expression within the dental follicle corresponds to an area of lower SIX1 expression, suggesting that it may be actively repressing SIX1+EYA transcriptional activity at this location. Within the developing mandible, there was strong expression of SOBP subjacent to that of SIX1 within the periosteum of the developing mandibular bone (**Fig. 9F**) and surrounding Meckel’s cartilage (**Fig. 9E**). This section also highlights the asymmetrical expression of SOBP in the condensed ectomesenchyme surrounding the developing incisor in this more anterior section (**Fig. 9E, F**). These data highlight the complex spatial distribution of SIX1 and SOBP within the developing mandible and incisor, which are both derived from the NCCs of the branchial arch and highlight the biological importance of understanding the dynamics of SIX1+EYA transcriptional regulation during craniofacial development.

**Figure 9:**
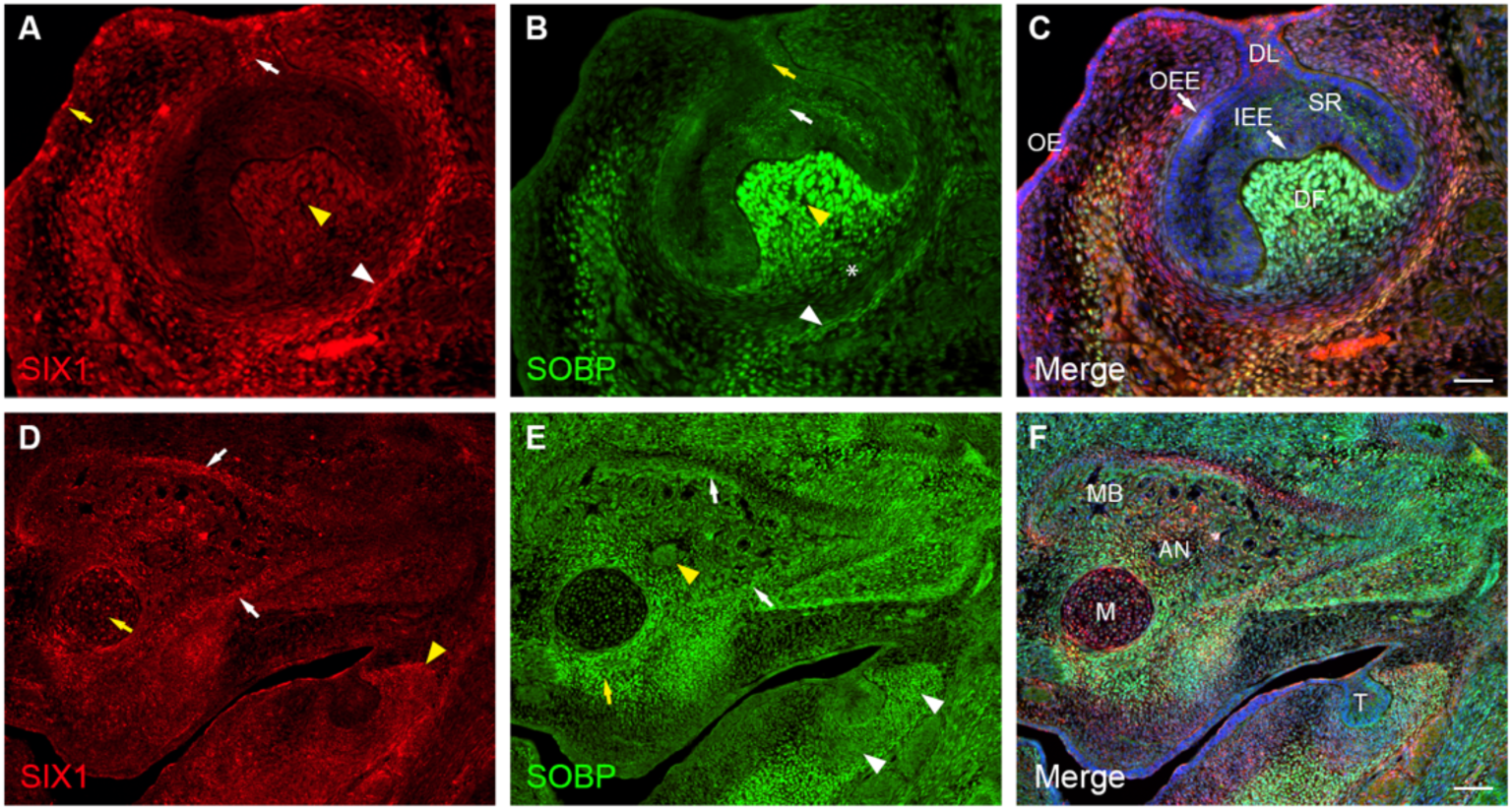
Transverse sections through the developing mouse mandible at E14.5 demonstrating the complex interactions between SIX1 and SOBP during both tooth and mandibular bone development. **A**) SIX1 is strongly expressed in the oral epithelium (OE, yellow arrow), the dental lamina (DL, white arrow) and in the border of the condensed ectomesenchyme surrounding the developing dental papilla (white arrowhead). It is also expressed, but at a lower level within the dental follicle (DF, yellow arrowhead). **B**) SOBP is weakly expressed within the DL (yellow arrow) and the stellate reticulum (white arrow), and this expression appears to be restricted to only one side of the developing tooth. SOBP is most strongly expressed within the DF (yellow arrowhead). There is then a region of low expression in the ectomesenchyme directly adjacent to the developing tooth (asterisks) and then there is SOBP expression within the boundary of the condensed ectomesenchyme (white arrowhead). **C**) This merged figure highlights that SOBP and SIX1 are co-expressed in the boundary of the condensed ectomesenchyme surrounding the developing tooth and that SOBP expression is much strongly than that of SIX1 within the DF. Scale bar is 50 μm. **D**) In the developing mandibular bone (MB), SIX1 is expressed within the periosteum (white arrow) and within Meckel’s cartilage (M, yellow arrow). There is also diffuse expression within the developing bone itself. The expression in the boundary of the condensed ectomesenchyme surrounding the developing tooth (T) also evident (yellow arrowhead). **E**) Interestingly, we observe strong expression of SOBP subjacent to that of SIX1 in the periosteum (white arrow). SOBP was also strongly expressed in the condensed ectomesenchyme surrounding Meckel’s cartilage (yellow arrow). Interestingly, there was also expression of SOBP within the developing alveolar nerve (AN, yellow arrowhead). This section also highlights the gradient of SOBP expression within the developing tooth (white arrowheads). **F**) This merged image highlights the complex spatial organization of SIX1 and SOBP, suggesting a dynamic regulation of SIX1+EYA transcriptional activity during mandibular development. Scale bar is 100 μm.

## Discussion

Branchio-oto-renal syndrome presents with variable craniofacial malformation that primarily affect the derivatives of the otic placode and mandibular arch (Smith, 1993, Moody et al., 2015, Ozaki et al., 2004, Laclef et al., 2003, Tavares et al., 2017, Guo et al., 2011, Pandur and Moody, 2000). To date, the most studied of these craniofacial phenotypes are those pertaining to the inner ear (derived from the otic placode) as undiagnosed hearing loss has long term effects on human development and is one of the characteristics of BOR (Smith, 1993, Moody et al., 2015, Neilson et al., 2010, Pandur and Moody, 2000, Neilson et al., 2020, Brugmann et al., 2004, Neilson et al., 2017, Keer et al., 2022, Tavares et al., 2021, Jourdeuil et al., 2023). Another possible cause of hearing loss in patients with BOR, which has received far less attention, is abnormalities in the middle ear ossicles which are derived from the ectomesenchyme of the mandibular arch (Smith, 1993, Moody et al., 2015, Tavares et al., 2017, Guo et al., 2011). As half of all patients with BOR present with mutations in either *SIX1* or its co-factor *EYA1*, previous research has focused on testing the role of *SIX1* and putative co-factors during otic placode development in different animal models (Neilson et al., 2020, Tavares et al., 2021, Neilson et al., 2017, Keer et al., 2022, Jourdeuil et al., 2023, Neilson et al., 2010, Moody et al., 2015). Work in *Xenopus* identified Pa2G4, Mcrs1, and Sobp as *bona fide* Six1 co-factors, that is, proteins that bind to Six1 and modulate its transcriptional activity (Keer et al., 2022, Neilson et al., 2017, Neilson et al., 2020, Tavares et al., 2021). Other than Mcrs1, the role of these co-factors has only been investigated during early inner ear development (Keer et al., 2022, Neilson et al., 2017, Neilson et al., 2020, Tavares et al., 2021) and their role in the NCC of the mandibular arch is largely unknown. Therefore, we set out to identify whether these proteins are co-expressed with SIX1 in the different domains of the mouse mandibular arch that will give rise to craniofacial structures altered in BOR patients. In addition, we investigated whether these proteins are *bona fide* SIX1 co-factors in the mouse and how they interact with the SIX1+EYA transcriptional complex to elucidate their role in the patterning of the mandibular arch ectomesenchyme.

### PA2G4 indirectly enhances SIX1+EYA transcriptional activity by facilitating translocation of EYA to the nucleus

PA2G4, also known as EBP1, can be differentially spliced to produce either a short PA2G4 (PA2G4-P42) or a long PA2G4 (PA2G4-P48) (Stevenson et al., 2020). The long PA2G4 (-P48), which is localized in both the cytosol and the nucleolus, is the predominant form of PA2G4 found in mammalian cells and has been shown to promote tumorigenesis (Stevenson et al., 2020). The short PA2G4 (-P42), which acts as a tumor suppressor, is expressed at relatively low levels in mammalian cells, and is located only in the cytosol (Stevenson et al., 2020, Squatrito et al., 2004). Pa2G4 was identified as a *bona fide* Six1 co-factor in *Xenopus* in 2017, at which time it was determined that this protein repressed the activity of two different reporters of Six1 transcriptional activity in human cells but enhanced the activity of these same reporters in *Xenopus* fibroblast-like cells (Neilson et al., 2017). The authors showed that Pa2G4 is expressed in the developing otic vesicle and branchial arches of *Xenopus* larvae, however, their work focused on inner ear development (Neilson et al., 2017). In agreement with the *Xenopus* data, our data demonstrate that PA2G4 is expressed throughout the mandibular arch at E10.5, extending from the proximal domain to the distal domain. Because SIX1 is required for proper upper and lower jaw development (Ozaki et al., 2004, Laclef et al., 2003, Tavares et al., 2017, Guo et al., 2011), this pattern of co-expression suggests that PA2G4 could regulate SIX1 function during the development of all these craniofacial elements. Further work using *Pa2G4* knockout mice is required to clarify whether this protein is required for the development of these structures and whether there are genetic interactions with *Six1*.

Within the cell, the expression of PA2G4-P48 only overlaps with SIX1 in the nucleolus and, in the mouse, PA2G4-P48 does not bind to SIX1 (in contrast to what has been observed in *Xenopus* (Neilson et al., 2017)) or EYA proteins. Interestingly, despite not binding, we determined that PA2G4-P48 is able to enhance the transcriptional activity of SIX1 when in the presence of EYA factors. Thus, the modulation of the SIX1+EYA transcriptional activity is likely being achieved by a protein intermediate. Indirect modulation is not surprisingly as binding to a transcription factor is not required for transcriptional regulation (Engeland, 2018). This has been well characterized through the indirect modulation of p53, whereby p53 upregulates the transcription of *p21*/*CDKN1A* (reviewed in (Engeland, 2018)). p21/CDKN1A then alters the post-translational modifications of components of the DREAM transcriptional complex, leading to either transcriptional repression or activation based on the proteins in the DREAM complex (reviewed in (Engeland, 2018)). Interestingly, PA2G4 has been shown to bind to the p53 E3 ligase HDM2, which can polyubiquinate p53 and lead to its degradation (Kim et al., 2010, Stevenson et al., 2020). It has also been proposed that PA2G4 can modulate p53 levels by stabilizing the HDM2 protein, which facilitates the interaction between HDM2 and Akt (Kim et al., 2012, Stevenson et al., 2020). Our data have potentially uncovered a similar mechanism of indirect modulation of SIX1+EYA transcriptional activity via PA2G4-mediated stabilization of either a part of the SIX1+EYA transcriptional complex or by stabilizing the interactions between EYA and an unknown factor in the cytosol, resulting in the translocation of EYA to the nucleus, making it available to SIX1 for transcriptional activation and induction of several genes including *Pa2G4* itself (**Fig. 10A**). Further work is required to identify whether this unknown intermediary protein is involved in mediating the interactions between PA2G4 and the members of the SIX1+EYA transcriptional complex.

**Figure 10.**
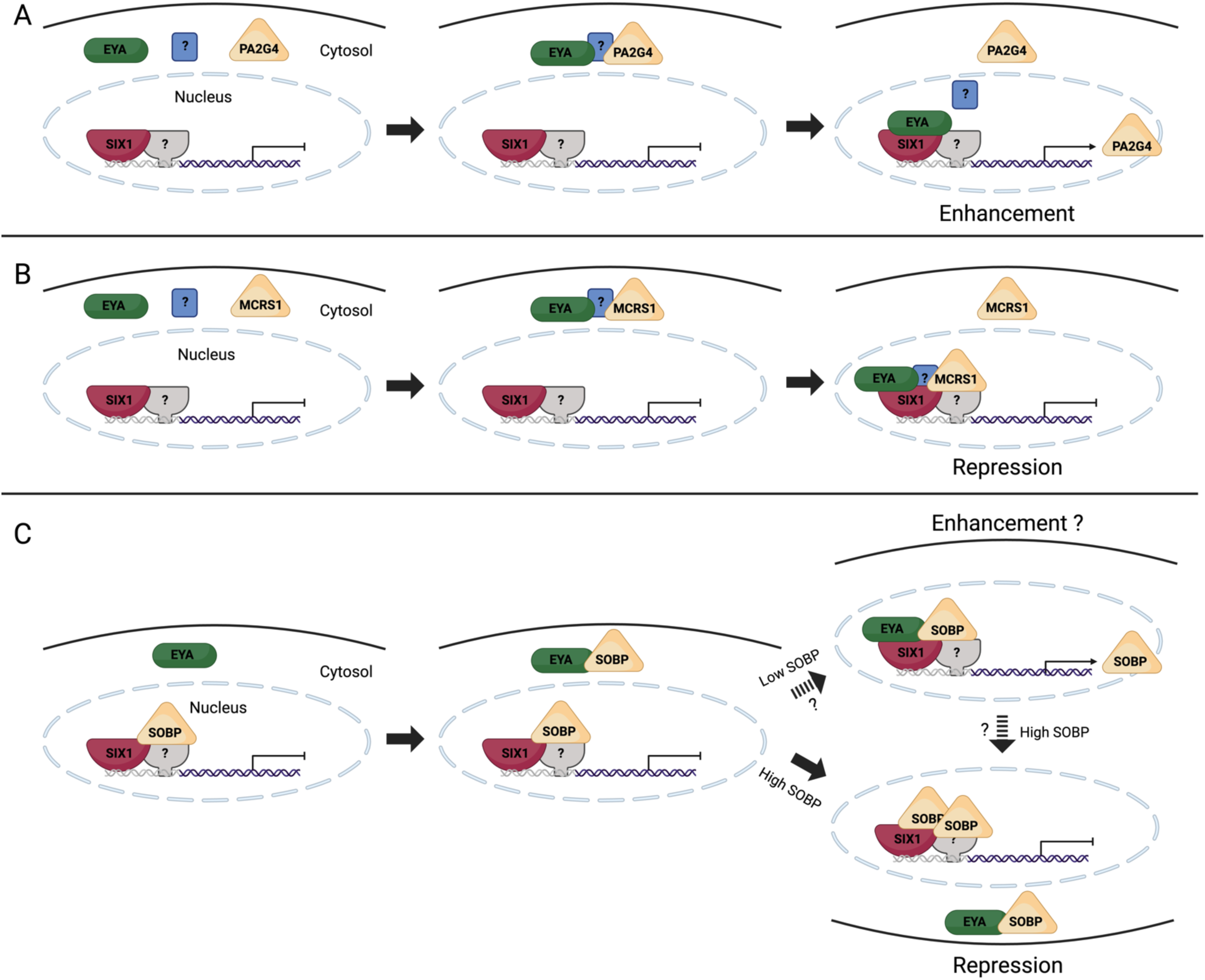
– Potential mechanisms regulating SIX1+EYA transcriptional activity via translocation of EYA to/from the nucleus. **A)** PA2G4 is mainly a cytosolic protein and does not bind to SIX1 or EYA. PA2G4 appears to act by stabilizing either a part of the SIX1+EYA transcriptional complex or the interactions between EYA and an unknown factor (?) in the cytosol. This results in the translocation of both EYA and PA2G4 to the nucleus, making it available to SIX1 for transcriptional activation and induction of several genes including *Pa2G4*. **B)** MCRS1 is mainly a cytosolic protein and binds to SIX1 but does not bind to EYA. MCRS1 indirectly facilitates translocation of EYA to the nucleus by an unknown factor (?). Repression of SIX1+EYA transcriptional activity is achieved by an unknown mechanism as EYA remains bound to SIX1 even in the presence of higher amounts of MCRS1 (Neilson et al., 2020). **C)** SOBP is a nuclear protein and binds to SIX1 and EYA. At lower amounts of SOBP and equimolar amounts of SIX1, EYA and SOBP, SOBP appears to be translocated to the cytosol while EYA is translocated to the nucleus. Repression of SIX1+EYA transcriptional activity is achieved by increasing amounts of SOBP leading to EYA translocation to the cytosol (Tavares et al., 2021). It is unknown if lower amounts of SOBP results in enhancement of the SIX1+EYA transcriptional activity which in turn would increase the levels of *Sobp* leading to repression.

### MCRS1 represses SIX1+EYA transcriptional activity and is able to translocate EYA to the nucleus

MCRS1, also known as MSP58, contributes to a number of cellular processes including regulation of various transcription factors, oncogenesis, and mediating the p53/p21 pathway via BRG1 (Yang et al., 2015a, Hsu et al., 2012, Xu et al., 2007). It has also been shown to relieve the transcriptional repression of the protein Daxx by binding to it and sequestering it in the nucleolus (Yang et al., 2015a, Lin and Shih, 2002). MCRS1 contains both a nuclear localization signal (NLS) and a nucleolar localization signal (NoLS) and its translocation to the nucleus is mediated through binding to importin α1 and α6 (Yang et al., 2015a). Interestingly, the nuclear translocation of MCRS1 is required for it to repress the expression of both p53 and p21, demonstrating its ability to control both gene expression and cell proliferation (Yang et al., 2015a, Hsu et al., 2012). Mcrs1 was identified as a *bona fide* Six1 co-factor in *Xenopus* in 2020, at which time it was shown that this protein binds to Six1 in both cultured cells and embryonic ectoderm and represses Six1+Eya1 transcriptional activity (Neilson et al., 2020). The authors further showed that loss of Mcrs1 resulted in the expansion of early neural plate and NC marker genes and a later reduction in otic vesicle gene expression (Neilson et al., 2020). A more recent study identified that Mcrs1 is required for inner ear formation and patterning and that loss of Mcrs1 resulted in defect NCC migration and abnormalities in the NC-derived craniofacial cartilages including the infrarostral, ceratohyal, otic capsule, and Meckel’s cartilages (Keer et al., 2022). Our data showed that, in mouse, MCRS1 is also a *bona fide* SIX1 co-factor. It is co-expressed with SIX1 within the mandibular arch at E10.5. Interestingly, the expression of MCRS1 also largely overlapped with that of PA2G4 within the arch at this stage. This pattern of co-expression suggests that there is likely a fine balance between repression and enhancement of SIX1+EYA transcriptional activity during the patterning of the mandibular arch ectomesenchyme. It will be important in future work to describe the phenotype, if any, of *Mcrs1* knockout mice and determine whether there are any interactions between *Pa2G4* and *Mcrs1*.

Within the cell, MCRS1 is primarily expressed in the cytosol but in the presence of SIX1, it becomes largely nuclear. Interestingly, a similar translocation was observed in the presence of both EYA1 and EYA2 without the addition of SIX1. We also noted that the translocation of MCRS1 to the nucleus was more pronounced with EYA2 than with EYA1. The translocation of MCRS1 to the nucleus in the presence of SIX1 was not entirely unexpected as we showed that in the mouse, as in *Xenopus*, MCRS1 binds to SIX1 (Neilson et al., 2020). We were surprised, however, that MCRS1 and EYA were translocated to the nucleus when co-transfected as our MCRS1 Co-IP data did not show any binding between MCRS1 and EYA. This suggests that MCRS1 may be stabilizing the interaction between EYA and another unknown protein to facilitate its translocation to the nucleus or binding directly to this intermediate protein to translocate EYA to the nucleus (**Fig. 10B**). It is possible that this unknown protein may be an importin, such as importin α1 or α6, as they are required for the nuclear translocation of MCRS1 to the nucleus (Yang et al., 2015a). Further, we noted that more MCRS1 was present in the nucleus than the cytosol when equimolar amounts of EYA and MCRS1 were transfected into MC3T3-E1 cells then in the same experiments with MCRS1 and SIX1. This finding was particularly interesting as, in *Xenopus*, while Sobp represses Six1+Eya1 transcriptional activity by competing with Eya1 for binding to Six1, resulting in the translocation of Eya1 to the cytosol (Tavares et al., 2021), Mcrs1 represses the transcriptional activity by an unknown mechanism that does not displace Eya1 from the Six1 transcriptional complex (Neilson et al., 2020). It is possible that MCRS1 is serving a similar role in the mouse, where at lower levels of SIX1, it acts as an enhancer and assists in the translocation of EYA to the nucleus but at higher levels of SIX1 binds to the SIX1+EYA transcriptional complex to repress transcription, although this has not yet been shown experimentally (**Fig. 10B**). The way in which MCRS1 indirectly translocates EYA to the nucleus but then represses SIX1+EYA transcriptional activity will require further study but may provide a mechanism for embryo to fine-tune gene expression of the downstream targets of SIX1 transcription during craniofacial development.

### SOBP function appears to be conserved in both mouse and *Xenopus*

*Sobp* was identified as the spontaneous recessive variant that causes deafness and vestibular-mediated circling behavior in the Jackson circler (*jc*) mouse (Chen et al., 2008, Calderon et al., 2006) and has been linked to intellectual disability, anterior maxillary, and mild hearing loss in humans (MRAMS; OMIM #613671). Sobp was identified as a *bona fide* Six1 co-factor that is required for the development of the inner ear in *Xenopus* (Tavares et al., 2021).

Interestingly, in *Xenopus*, Sobp is only expressed in the developing inner ear and did not appear to be expressed in the branchial arches (Tavares et al., 2021, Neilson et al., 2010). Our findings in mouse identified that SOBP is expressed in a very restricted area of ectoderm and ectomesenchyme of the mandibular arch in the oral half, approximately halfway between the proximal and distal domains of the arch. Based on its positional location, it is likely that this corresponds to the developing odontogenic mesenchyme and dental lamina at E10.5 (Sharpe, 2001, Tucker and Sharpe, 2004) and could reflect a novel function for SOBP in mammals. Our results demonstrate that at E14.5, during incisor development, there is an asymmetric gradient of SOBP within the dental lamina and stellate reticulum but that it is absent from the oral epithelium. We further showed that SOBP is strongly expressed in the dental follicle, where we noted lower expression of SIX1, suggesting an important role for SOBP in regulating incisor development, a novel function for SOBP in odontogenesis. These findings present exciting avenues of future study into the role of SOBP regulation of SIX1 during odontogenesis, especially given that SIX1 has recently been shown to be required for lower incisor development (Takahashi et al., 2020).

As SOBP is a nuclear protein, co-expression of SIX1 and SOBP did not change the localization of either protein. However, we found that when SOBP and EYA1 were co-transfected, the expression of each became punctate and that EYA1 was partially translocated to the nucleus while SOBP was partially translocated to the cytosol. It was previously described in *Xenopus* that lower levels of Sobp stabilize the interaction between Six1+Eya1 while higher levels of Sobp disrupt this interaction by competing with Eya1 for binding to Six1 and translocating Eya1 from the nucleus to the cytosol, resulting in transcriptional repression (Tavares et al., 2021). Our results demonstrate a similar dual action of SOBP, whereby it is both able to directly bind to EYA and translocate it to the nucleus at lower levels of SIX1 (*i.e.*, in this case, the endogenously expressed SIX1 in the MC3T3-E1 cells) but at higher levels (*i.e.,* equimolar amounts of SOBP, EYA, and SIX1) represses transcriptional activity of the SIX1+EYA complex. It is possible that this constitutes a feedback loop wherein at low SOBP levels, EYA is translocated to the nucleus; however, when there is too much SOBP, it then removes EYA from the nucleus and represses SIX1 transcriptional activity. It is presently unknown whether lower amounts of SOBP would result in enhanced SIX1 transcriptional activity or whether expression of SIX1-regulated genes, including *Sobp*, would in turn increase the expression of this co-repressor creating a negative feedback loop (**Fig. 10C**). Interestingly, our expression data revealed that the expression pattern of SIX1 and SOBP in the developing mandible and incisor is similar to the previously reported pattern in the pre-placodal ectoderm and otic vesicle (Tavares et al., 2021). In these two embryonic structures, *Sobp* and *Six1* are co-expressed but their expression appears inverted, with higher levels of *Sobp* corresponding to lower levels of *Six1* expression and vice versa. This expression pattern suggests a conserved mechanism for SIX1+EYA transcriptional activity modulation by SOBP within very fine parameters during ontogeny in specific tissues. This exciting finding represents fertile ground for further study.

## Conclusion

This study highlights the complex mechanisms regulating SIX1+EYA transcriptional activity and suggests there are multiple, potentially redundant, mechanisms within the mouse to control fate determination via SIX1 downstream targets. By identifying other putative SIX1 co-factors and the precise levels of SIX1 required for fate determination, we believe it will be possible to further unravel the role of SIX1 during craniofacial development, especially the elements most affected in patients with BOR. This is also an especially exciting avenue of further study as there is now evidence that the mechanisms controlling specific levels of SIX1 may be affected in patients presenting with craniosynostoses.

## References

Alexander, C., Zuniga, E., Blitz, I. L., Wada, N., Le Pabic, P., Javidan, Y., Zhang, T., Cho, K. W., Crump, J. G. & Schilling, T. F. 2011. Combinatorial roles for BMPs and Endothelin 1 in patterning the dorsal-ventral axis of the craniofacial skeleton. Development, 138, 5135–46.

Barske, L., Askary, A., Zuniga, E., Balczerski, B., Bump, P., Nichols, J. T. & Crump, J. G. 2016. Competition between Jagged-Notch and Endothelin1 Signaling Selectively Restricts Cartilage Formation in the Zebrafish Upper Face. PLoS Genet, 12, e1005967.

Baxi, A., Jourdeuil, K., Cox, T. C., Clouthier, D. E. & Tavares, A. L. P. 2023. Transcriptomic analysis reveals the role of SIX1 in mouse cranial neural crest patterning and bone development. Dev Dyn, 252, 1303–1315.

Brugmann, S. A., Pandur, P. D., Kenyon, K. L., Pignoni, F. & Moody, S. A. 2004. Six1 promotes a placodal fate within the lateral neurogenic ectoderm by functioning as both a transcriptional activator and repressor. Development, 131, 5871–81.

Buller, C., Xu, X., Marquis, V., Schwanke, R. & Xu, P. X. 2001. Molecular effects of Eya1 domain mutations causing organ defects in BOR syndrome. Hum Mol Genet, 10, 2775–81.

Calderon, A., Derr, A., Stagner, B. B., Johnson, K. R., Martin, G. & NOBEN-Trauth, K. 2006. Cochlear developmental defect and background-dependent hearing thresholds in the Jackson circler (jc) mutant mouse. Hear Res, 221, 44–58.

Calpena, E., Wurmser, M., Mcgowan, S. J., Atique, R., Bertola, D. R., Cunningham, M. L., Gustafson, J. A., Johnson, D., Morton, J. E. V., PASSOS-Bueno, M. R., Timberlake, A. T., Lifton, R. P., Wall, S. A., Twigg, S. R. F., Maire, P. & Wilkie, A. O. M. 2022. Unexpected role of SIX1 variants in craniosynostosis: expanding the phenotype of SIX1-related disorders. J Med Genet, 59, 165–169.

Chen, Z., Montcouquiol, M., Calderon, R., Jenkins, N. A., Copeland, N. G., Kelley, M. W. & NOBEN-Trauth, K. 2008. Jxc1/Sobp, Encoding a Nuclear Zinc Finger Protein, Is Critical for Cochlear Growth, Cell Fate, and Patterning of the Organ of Corti. The Journal of Neuroscience, 28, 6633–6641.

Clouthier, D. E., Hosoda, K., Richardson, J. A., Williams, S. C., Yanagisawa, H., Kuwaki, T., Kumada, M., Hammer, R. E. & Yanagisawa, M. 1998. Cranial and cardiac neural crest defects in endothelin-A receptor-deficient mice. Development, 125, 813–24.

Clouthier, D. E., Williams, S. C., Yanagisawa, H., Wieduwilt, M., Richardson, J. A. & Yanagisawa, M. 2000. Signaling pathways crucial for craniofacial development revealed by endothelin-A receptor-deficient mice. Dev Biol, 217, 10–24.

Du, X., Wang, Q., Hirohashi, Y. & Greene, M. I. 2006. Dipa, which can localize to the centrosome, associates with p78/MCRS1/MSP58 and acts as a repressor of gene transcription. Exp Mol Pathol, 81, 184–90.

Engeland, K. 2018. Cell cycle arrest through indirect transcriptional repression by p53: I have a DREAM. Cell Death Differ, 25, 114–132.

Figeac, N., Serralbo, O., Marcelle, C. & Zammit, P. S. 2014. ErbB3 binding protein-1 (Ebp1) controls proliferation and myogenic differentiation of muscle stem cells. Dev Biol, 386, 135–51.

Ford, H. L., Landesman-Bollag, E., Dacwag, C. S., Stukenberg, P. T., Pardee, A. B. & Seldin, D. C. 2000. Cell cycle-regulated phosphorylation of the human SIX1 homeodomain protein. J Biol Chem, 275, 22245–54.

Franz-Odendaal, T. A., Hall, B. K. & Witten, P. E. 2006. Buried alive: how osteoblasts become osteocytes. Dev Dyn, 235, 176–90.

Guo, C., Sun, Y., Zhou, B., Adam, R. M., Li, X., Pu, W. T., Morrow, B. E., Moon, A. & Li, X. 2011. A Tbx1-Six1/Eya1-Fgf8 genetic pathway controls mammalian cardiovascular and craniofacial morphogenesis. J Clin Invest, 121, 1585–95.

Hsu, C. C., Lee, Y. C., Yeh, S. H., Chen, C. H., Wu, C. C., Wang, T. Y., Chen, Y. N., Hung, L. Y., Liu, Y. W., Chen, H. K., Hsiao, Y. T., Wang, W. S., Tsou, J. H., Tsou, Y. H., Wu, M. H., Chang, W. C. & Lin, D. Y. 2012. 58-kDa microspherule protein (MSP58) is novel Brahma-related gene 1 (BRG1)-associated protein that modulates p53/p21 senescence pathway. J Biol Chem, 287, 22533–48.

Ikeda, K., Watanabe, Y., Ohto, H. & Kawakami, K. 2002. Molecular interaction and synergistic activation of a promoter by Six, Eya, and Dach proteins mediated through CREB binding protein. Mol Cell Biol, 22, 6759–66.

Ivanova, A. V., Ivanov, S. V. & Lerman, M. L. 2005. Association, mutual stabilization, and transcriptional activity of the STRA13 and MSP58 proteins. Cell Mol Life Sci, 62, 471–84.

Jourdeuil, K., Neilson, K. M., Cousin, H., Tavares, A. L. P., Majumdar, H. D., Alfandari, D. & Moody, S. A. 2023. Zmym4 is required for early cranial gene expression and craniofacial cartilage formation. Front Cell Dev Biol, 11, 1274788.

Kaufman, M. H. 1994. The Atlas of Mouse Development, Oxford, Uk, Elsevier Academic Press.

Keer, S., Cousin, H., Jourdeuil, K., Neilson, K. M., Tavares, A. L. P., Alfandari, D. & Moody, S. A. 2022. Mcrs1 is required for branchial arch and cranial cartilage development. Dev Biol, 489, 62–75.

Kenyon, K. L., Li, D. J., Clouser, C., Tran, S. & Pignoni, F. 2005a. Fly SIX-type homeodomain proteins Sine oculis and Optix partner with different cofactors during eye development. Dev Dyn, 234, 497–504.

Kenyon, K. L., YANG-Zhou, D., Cai, C. Q., Tran, S., Clouser, C., Decene, G., Ranade, S. & Pignoni, F. 2005b. Partner specificity is essential for proper function of the SIX-type homeodomain proteins Sine oculis and Optix during fly eye development. Dev Biol, 286, 158–68.

Kim, C. K., Lee, S. B., Nguyen, T. L., Lee, K. H., Um, S. H., Kim, J. & Ahn, J. Y. 2012. Long isoform of ErbB3 binding protein, p48, mediates protein kinase B/Akt-dependent HDM2 stabilization and nuclear localization. Exp Cell Res, 318, 136-43.

Kim, C. K., Nguyen, T. L., Joo, K. M., Nam, D. H., Park, J., Lee, K. H., Cho, S. W. & Ahn, J. Y. 2010. Negative regulation of p53 by the long isoform of ErbB3 binding protein Ebp1 in brain tumors. Cancer Res, 70, 9730–41.

Klingbeil, K. D., Greenland, C. M., Arslan, S., LLAMOS Paneque, A., Gurkan, H., DEMIR Ulusal, S., Maroofian, R., Carrera-Gonzalez, A., MONTUFAR-Armendariz, S., Paredes, R., Elcioglu, N., Menendez, I., Behnam, M., Foster, J., 2nd, Guo, S., Escarfuller, S., Cengiz, F. B., Duman, D., Bademci, G. & Tekin, M. 2017. Novel EYA1 variants causing Branchio-oto-renal syndrome. Int J Pediatr Otorhinolaryngol, 98, 59-63.

Ko, H. R., Chang, Y. S., Park, W. S. & Ahn, J. Y. 2016. Opposing roles of the two isoforms of ErbB3 binding protein 1 in human cancer cells. Int J Cancer, 139, 1202–8.

Kobayashi, M., Nishikawa, K., Suzuki, T. & Yamamoto, M. 2001. The homeobox protein Six3 interacts with the Groucho corepressor and acts as a transcriptional repressor in eye and forebrain formation. Dev Biol, 232, 315–26.

Kochhar, A., Fischer, S. M., Kimberling, W. J. & Smith, R. J. 2007. Branchio-oto-renal syndrome. Am J Med Genet A, 143a, 1671-8.

Kowalinski, E., Bange, G., Bradatsch, B., Hurt, E., Wild, K. & Sinning, I. 2007. The crystal structure of Ebp1 reveals a methionine aminopeptidase fold as binding platform for multiple interactions. FEBS Lett, 581, 4450–4.

Laclef, C., Souil, E., Demignon, J. & Maire, P. 2003. Thymus, kidney and craniofacial abnormalities in Six 1 deficient mice. Mech Dev, 120, 669–79.

Le Lievre, C. S. 1978. Participation of neural crest-derived cells in the genesis of the skull in birds. J Embryol Exp Morphol, 47, 17–37.

Le Lièvre, C. S. & Le Douarin, N. M. 1975. Mesenchymal derivatives of the neural crest: analysis of chimaeric quail and chick embryos. Journal of Embryology and Experimental Morphology, 34, 125–154.

Lee, K. Y., Kim, S., Kim, U. K., Ki, C. S. & Lee, S. H. 2007. Novel EYA1 mutation in a Korean branchio-oto-renal syndrome family. Int J Pediatr Otorhinolaryngol, 71, 169–74.

Li, J., Lee, G. I., VAN Doren, S. R. & Walker, J. C. 2000. The FHA domain mediates phosphoprotein interactions. J Cell Sci, 113 Pt 23, 4143–9.

Li, X., Oghi, K. A., Zhang, J., Krones, A., Bush, K. T., Glass, C. K., Nigam, S. K., Aggarwal, A. K., Maas, R., Rose, D. W. & Rosenfeld, M. G. 2003. Eya protein phosphatase activity regulates Six1-Dach-Eya transcriptional effects in mammalian organogenesis. Nature, 426, 247–54.

Li, Y., Manaligod, J. M. & Weeks, D. L. 2010. EYA1 mutations associated with the brancio-oto-renal syndrome result in defective otic development in Xenopus laevis. Biology of the Cell, 102, 277–292.

Lin, D. Y. & Shih, H. M. 2002. Essential role of the 58-kDa microspherule protein in the modulation of Daxx-dependent transcriptional repression as revealed by nucleolar sequestration. J Biol Chem, 277, 25446–56.

Lu, Y., Zhou, H., Chen, W., Zhang, Y. & Hamburger, A. W. 2011. The ErbB3 binding protein EBP1 regulates ErbB2 protein levels and tamoxifen sensitivity in breast cancer cells. Breast Cancer Res Treat, 126, 27–36.

Monie, T. P., Perrin, A. J., Birtley, J. R., Sweeney, T. R., Karakasiliotis, I., Chaudhry, Y., Roberts, L. O., Matthews, S., Goodfellow, I. G. & Curry, S. 2007. Structural insights into the transcriptional and translational roles of Ebp1. EMBO J, 26, 3936–44.

Moody, S. A. & Lamantia, A. S. 2015. Transcriptional regulation of cranial sensory placode development. Curr Top Dev Biol, 111, 301–50.

Moody, S. A., Neilson, K. M., Kenyon, K. L., Alfandari, D. & Pignoni, F. 2015. Using Xenopus to discover new genes involved in branchiootorenal spectrum disorders. Comp Biochem Physiol C Toxicol Pharmacol, 178, 16–24.

Neilson, K. M., Abbruzzesse, G., Kenyon, K., Bartolo, V., Krohn, P., Alfandari, D. & Moody, S. A. 2017. Pa2G4 is a novel Six1 co-factor that is required for neural crest and otic development. Dev Biol, 421, 171–182.

Neilson, K. M., Keer, S., Bousquet, N., Macrorie, O., Majumdar, H. D., Kenyon, K. L., Alfandari, D. & Moody, S. A. 2020. Mcrs1 interacts with Six1 to influence early craniofacial and otic development. Dev Biol, 467, 39–50.

Neilson, K. M., Pignoni, F., Yan, B. & Moody, S. A. 2010. Developmental expression patterns of candidate cofactors for vertebrate six family transcription factors. Dev Dyn, 239, 3446–66.

Ohto, H., Kamada, S., Tago, K., Tominaga, S. I., Ozaki, H., Sato, S. & Kawakami, K. 1999. Cooperation of six and eya in activation of their target genes through nuclear translocation of Eya. Mol Cell Biol, 19, 6815–24.

Ozaki, H., Nakamura, K., Funahashi, J., Ikeda, K., Yamada, G., Tokano, H., Okamura, H. O., Kitamura, K., Muto, S., Kotaki, H., Sudo, K., Horai, R., Iwakura, Y. & Kawakami, K. 2004. Six1 controls patterning of the mouse otic vesicle. Development, 131, 551–62.

Pandur, P. D. & Moody, S. A. 2000. Xenopus Six1 gene is expressed in neurogenic cranial placodes and maintained in the differentiating lateral lines. Mech Dev, 96, 253–7.

Patrick, A. N., Cabrera, J. H., Smith, A. L., Chen, X. S., Ford, H. L. & Zhao, R. 2013. Structure-function analyses of the human SIX1-EYA2 complex reveal insights into metastasis and BOR syndrome. Nat Struct Mol Biol, 20, 447–53.

Patrick, A. N., Schiemann, B. J., Yang, K., Zhao, R. & Ford, H. L. 2009. Biochemical and functional characterization of six SIX1 Branchio-oto-renal syndrome mutations. J Biol Chem, 284, 20781–90.

Pritchard, A. B., Kanai, S. M., Krock, B., Schindewolf, E., OLIVER- Krasinski, J., Khalek, N., Okashah, N., Lambert, N. A., Tavares, A. L. P., Zackai, E. & Clouthier, D. E. 2020. Loss-of-function of Endothelin receptor type A results in Oro-Oto-Cardiac syndrome. Am J Med Genet A, 182, 1104–1116.

Rafiq, A., Aashaq, S., Jan, I. & Beigh, M. A. 2021. SIX1 transcription factor: A review of cellular functions and regulatory dynamics. Int J Biol Macromol, 193, 1151–1164.

Raja, S. J., Charapitsa, I., Conrad, T., Vaquerizas, J. M., Gebhardt, P., Holz, H., Kadlec, J., Fraterman, S., Luscombe, N. M. & Akhtar, A. 2010. The nonspecific lethal complex is a transcriptional regulator in Drosophila. Mol Cell, 38, 827–41.

Ruf, R. G., Xu, P. X., Silvius, D., Otto, E. A., Beekmann, F., Muerb, U. T., Kumar, S., Neuhaus, T. J., Kemper, M. J., Raymond, R. M., JR., Brophy, P. D., Berkman, J., Gattas, M., Hyland, V., Ruf, E. M., Schwartz, C., Chang, E. H., Smith, R. J., Stratakis, C. A., Weil, D., Petit, C. & Hildebrandt, F. 2004. SIX1 mutations cause branchio-oto-renal syndrome by disruption of EYA1-SIX1-DNA complexes. Proc Natl Acad Sci U S A, 101, 8090–5.

Saint-Jeannet, J. P. & Moody, S. A. 2014. Establishing the pre-placodal region and breaking it into placodes with distinct identities. Dev Biol, 389, 13–27.

Sanggaard, K. M., Rendtorff, N. D., Kjaer, K. W., Eiberg, H., Johnsen, T., Gimsing, S., Dyrmose, J., Nielsen, K. O., Lage, K. & Tranebjaerg, L. 2007. Branchio-oto-renal syndrome: detection of EYA1 and SIX1 mutations in five out of six Danish families by combining linkage, MLPA and sequencing analyses. Eur J Hum Genet, 15, 1121–31.

Senel, E., Kocak, H., Akbiyik, F., Saylam, G., Gulleroglu, B. N. & Senel, S. 2009. From a branchial fistula to a branchiootorenal syndrome: a case report and review of the literature. J Pediatr Surg, 44, 623–5.

Seto, Y., Ogihara, R., Takizawa, K. & Eiraku, M. 2024. In vitro induction of patterned branchial arch-like aggregate from human pluripotent stem cells. Nat Commun, 15, 1351.

Shah, A. M., Krohn, P., Baxi, A. B., Tavares, A. L. P., Sullivan, C. H., Chillakuru, Y. R., Majumdar, H. D., Neilson, K. M. & Moody, S. A. 2020. Six1 proteins with human branchio-oto-renal mutations differentially affect cranial gene expression and otic development. Dis Model Mech, 13.

Sharpe, P. T. 2001. Neural crest and tooth morphogenesis. Adv Dent Res, 15, 4–7.

Sheikh, B. N., Guhathakurta, S. & Akhtar, A. 2019. The non-specific lethal (NSL) complex at the crossroads of transcriptional control and cellular homeostasis. EMBO Rep, 20, e47630.

Shimono, K., Shimono, Y., Shimokata, K., Ishiguro, N. & Takahashi, M. 2005. Microspherule protein 1, Mi-2beta, and RET finger protein associate in the nucleolus and up-regulate ribosomal gene transcription. J Biol Chem, 280, 39436-47.

Silver, S. J., Davies, E. L., Doyon, L. & Rebay, I. 2003. Functional dissection of eyes absent reveals new modes of regulation within the retinal determination gene network. Mol Cell Biol, 23, 5989–99.

Smith, R. J. H. 1993. Branchiootorenal Spectrum Disorder. In: Adam, M. P., Feldman, J., Mirzaa, G. M., Pagon, R. A., Wallace, S. E., Bean, L. J. H., Gripp, K. W. & Amemiya, A. (eds.) GeneReviews((R)). Seattle (WA).

Squatrito, M., Mancino, M., Donzelli, M., Areces, L. B. & Draetta, G. F. 2004. EBP1 is a nucleolar growth-regulating protein that is part of pre-ribosomal ribonucleoprotein complexes. Oncogene, 23, 4454–65.

Stevenson, B. W., Gorman, M. A., Koach, J., Cheung, B. B., Marshall, G. M., Parker, M. W. & Holien, J. K. 2020. A structural view of PA2G4 isoforms with opposing functions in cancer. J Biol Chem, 295, 16100–16112.

Takahashi, M., Ikeda, K., Ohmuraya, M., Nakagawa, Y., Sakuma, T., Yamamoto, T. & Kawakami, K. 2020. Six1 is required for signaling center formation and labial-lingual asymmetry in developing lower incisors. Dev Dyn, 249, 1098–1116.

Takahashi, Y., Bontoux, M. & Le Douarin, N. M. 1991. Epithelio--mesenchymal interactions are critical for Quox 7 expression and membrane bone differentiation in the neural crest derived mandibular mesenchyme. EMBO J, 10, 2387–93.

Tavares, A. L. & Clouthier, D. E. 2015. Cre recombinase-regulated Endothelin1 transgenic mouse lines: novel tools for analysis of embryonic and adult disorders. Dev Biol, 400, 191–201.

Tavares, A. L. P., Cox, T. C., Maxson, R. M., Ford, H. L. & Clouthier, D. E. 2017. Negative regulation of endothelin signaling by SIX1 is required for proper maxillary development. Development, 144, 2021–2031.

Tavares, A. L. P., Garcia, E. L., Kuhn, K., Woods, C. M., Williams, T. & Clouthier, D. E. 2012. Ectodermal-derived Endothelin1 is required for patterning the distal and intermediate domains of the mouse mandibular arch. Developmental Biology, 371, 47–56.

Tavares, A. L. P., Jourdeuil, K., Neilson, K. M., Majumdar, H. D. & Moody, S. A. 2021. Sobp modulates the transcriptional activation of Six1 target genes and is required during craniofacial development. Development, 148.

Teng, C. S., Yen, H. Y., Barske, L., Smith, B., Llamas, J., Segil, N., Go, J., SANCHEZ-Lara, P. A., Maxson, R. E. & Crump, J. G. 2017. Requirement for Jagged1-Notch2 signaling in patterning the bones of the mouse and human middle ear. Sci Rep, 7, 2497.

Tucker, A. & Sharpe, P. 2004. The cutting-edge of mammalian development; how the embryo makes teeth. Nat Rev Genet, 5, 499–508.

Tucker, A. S., Yamada, G., Grigoriou, M., Pachnis, V. & Sharpe, P. T. 1999. Fgf-8 determines rostral-caudal polarity in the first branchial arch. Development, 126, 51–61.

Wang, D., Christensen, K., Chawla, K., Xiao, G., Krebsbach, P. H. & Franceschi, R. T. 1999. Isolation and characterization of MC3T3-E1 preosteoblast subclones with distinct in vitro and in vivo differentiation/mineralization potential. J Bone Miner Res, 14, 893–903.

Wu, Z., Rao, Y., Zhang, S., Kim, E. J., Oki, S., Harada, H., Cheung, M. & Jung, H. S. 2019. Cis-control of Six1 expression in neural crest cells during craniofacial development. Dev Dyn, 248, 1264–1272.

Xia, X., Cheng, A., Lessor, T., Zhang, Y. & Hamburger, A. W. 2001. Ebp1, an ErbB-3 binding protein, interacts with Rb and affects Rb transcriptional regulation. J Cell Physiol, 187, 209–17.

Xu, P. X. 2013. The EYA-SO/SIX complex in development and disease. Pediatric Nephrology, 28, 843–854.

Xu, Y., Zhang, J. & Chen, X. 2007. The activity of p53 is differentially regulated by Brm- and Brg1-containing SWI/SNF chromatin remodeling complexes. J Biol Chem, 282, 37429–35.

Yang, C.-P., Chiang, C.-W., Chen, C.-H., Lee, Y.-C., We, M.-H., Tsou, Y.-H., Yang, Y.-S., Chang, W.-C. & Lin, D.-Y. 2015a. Identification and characterization of nuclear and nucleolar localization signals in 58-kDa microspherule protein (MSP58). Journal of Biomedical Science, 22, 33.

Yang, C. P., Chiang, C. W., Chen, C. H., Lee, Y. C., Wu, M. H., Tsou, Y. H., Yang, Y. S., Chang, W. C. & Lin, D. Y. 2015b. Identification and characterization of nuclear and nucleolar localization signals in 58-kDa microspherule protein (MSP58). J Biomed Sci, 22, 33.

Yoo, J. Y., Wang, X. W., Rishi, A. K., Lessor, T., Xia, X. M., Gustafson, T. A. & Hamburger, A. W. 2000. Interaction of the PA2G4 (EBP1) protein with ErbB-3 and regulation of this binding by heregulin. Br J Cancer, 82, 683–90.

Zhang, T., Xu, J. & Xu, P. X. 2021. Eya2 expression during mouse embryonic development revealed by Eya2(lacZ) knockin reporter and homozygous mice show mild hearing loss. Dev Dyn, 250, 1450–1462.

Zhang, Y., Lu, Y., Zhou, H., Lee, M., Liu, Z., Hassel, B. A. & Hamburger, A. W. 2008. Alterations in cell growth and signaling in ErbB3 binding protein-1 (Ebp1) deficient mice. BMC Cell Biol, 9, 69.

Zhou, H., MAZAN-Mamczarz, K., Martindale, J. L., Barker, A., Liu, Z., Gorospe, M., Leedman, P. J., Gartenhaus, R. B., Hamburger, A. W. & Zhang, Y. 2010. Post-transcriptional regulation of androgen receptor mRNA by an ErbB3 binding protein 1 in prostate cancer. Nucleic Acids Res, 38, 3619–31.

Zuniga, E., Rippen, M., Alexander, C., Schilling, T. F. & Crump, J. G. 2011. Gremlin 2 regulates distinct roles of BMP and Endothelin 1 signaling in dorsoventral patterning of the facial skeleton. Development, 138, 5147–56.

