## Supplementary figures for "Direct and Indirect Regulation of SIX1+EYA Transcriptional Activity by PA2G4, MCRS1, and SOBP"

*Supplementary Material*

### 1.1 Supplementary Figures

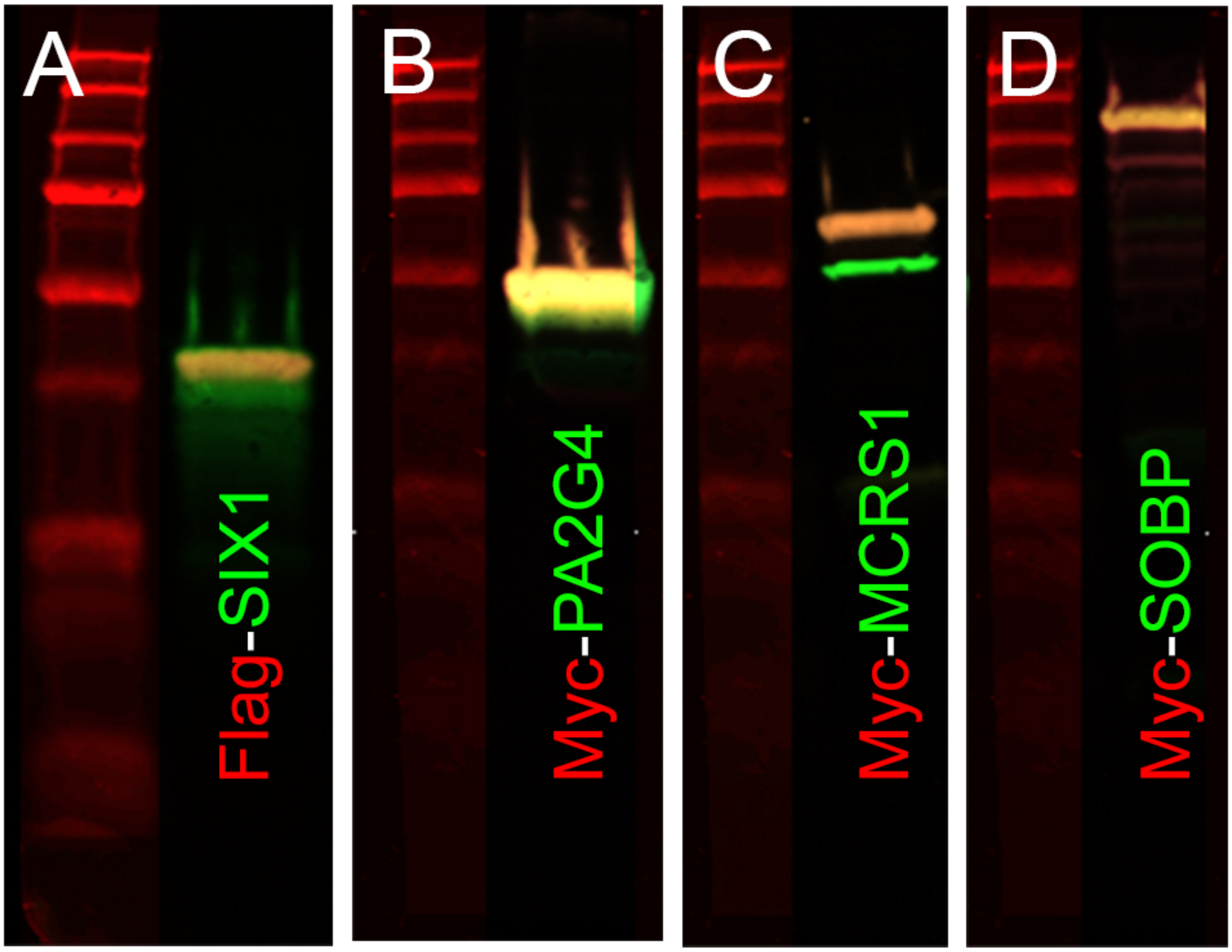


**Supplemental Figure 1: Antibody validation of SIX1, PA2G4, MCRS1, and SOBP.** Multiplex fluorescent Western blot detection. **A.** Flag and SIX1, **B.** Myc and PA2G4, **C.** Myc and MCRS1, **D.** Myc and SOBP. The expression of the tag (red) and the antibody for the protein itself (green) completely overlapped (yellow).

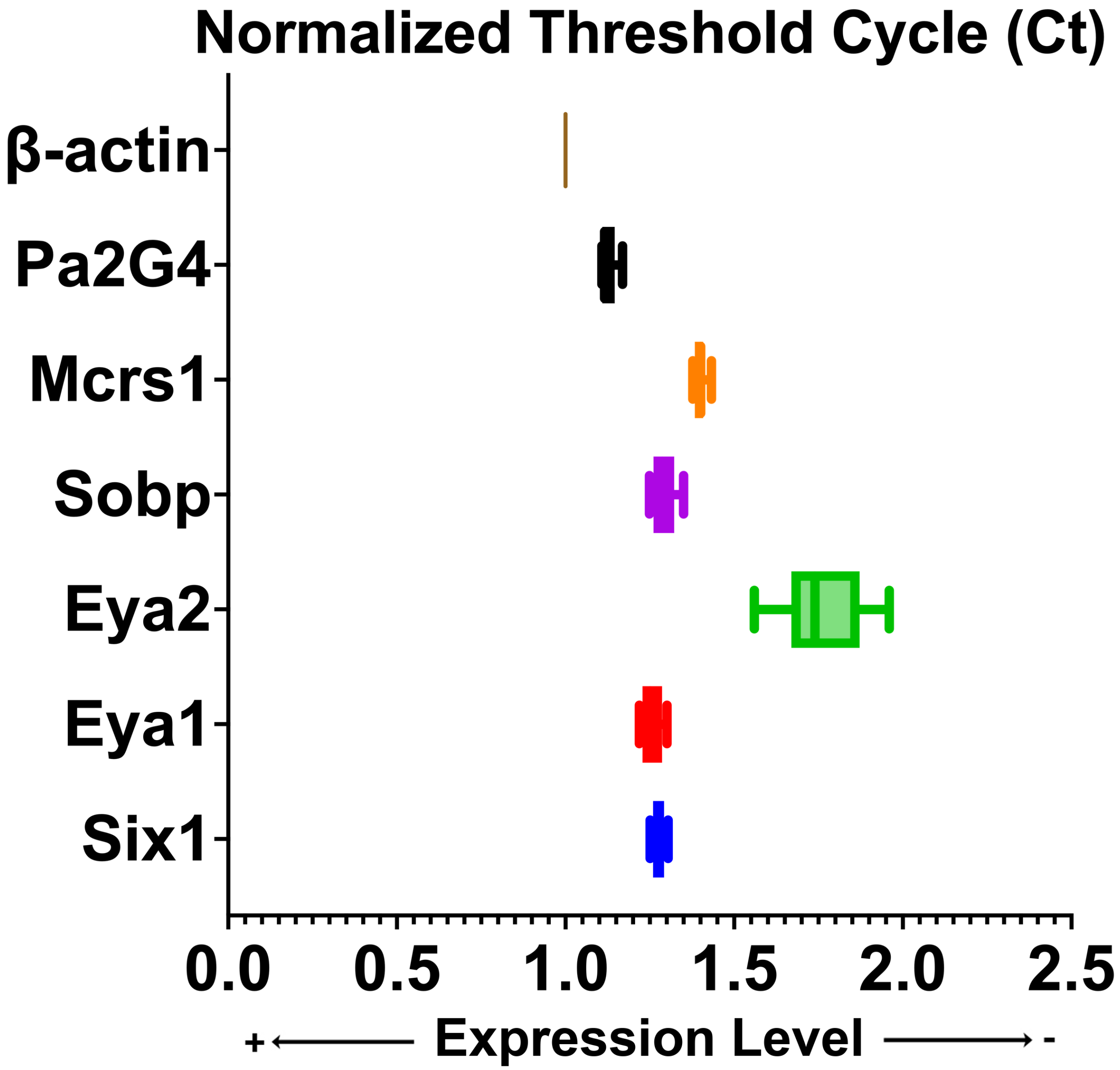


**Supplemental Figure 2: mRNA expression level analysis for *Six1*, *Eya1*, *Eya2*, *Pa2G4*, *Mcrs1* and *Sobp* in untransfected MC3T3-E1 cells by qPCR.** Normalized threshold cycles (Ct) for each analyzed gene were plotted relative to b-actin (*Actb*) Results show that *Six1*, *Eya1* and *Sobp* have similar expression levels. *Mcrs1* has a slightly lower expression level and *Pa2G4* has the highest expression level. *Eya2* does not seem to be expressed in these cells or if it is expressed, the very low expression level is not accurately amplified by qPCR.

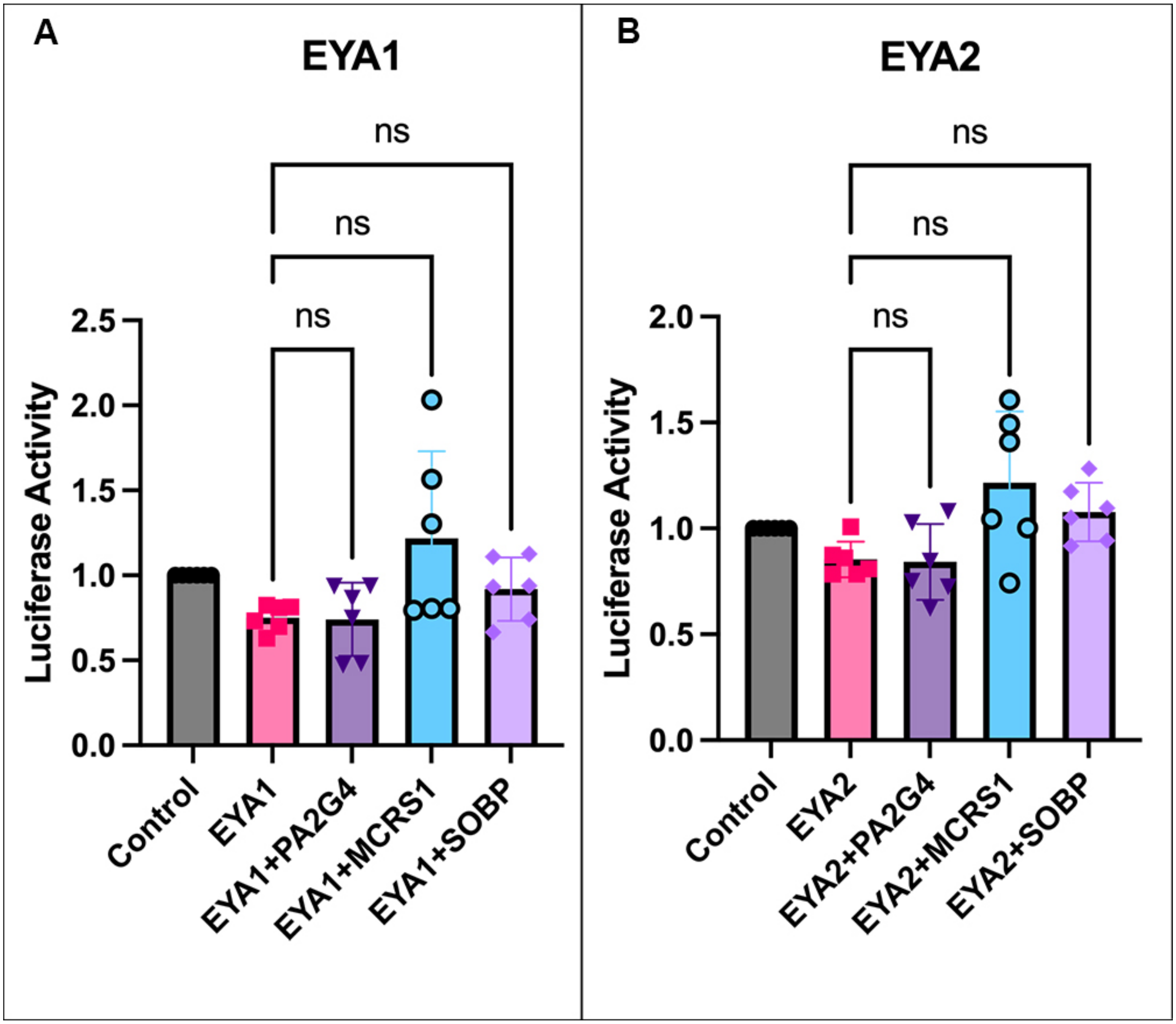


**Supplemental Figure 3: Neither *Pa2G4*, *Mcrs1*, nor *Sobp* significantly alter the luciferase activity of the reporter vector with *Eya1* or *Eya2* but without *Six1*.** (**A-B**) Graphs depicting the luciferase activity of the pGL3-6xMEF3-luciferase reporter in MC3T3-E1 cells transfected with different combinations of either control vector, HA-*Eya1*, HA-*Eya2*, Myc-*Pa2G4,* Myc-*Mcrs1*, and/or Myc-*Sobp*. Data were normalized to CMV-Renilla and are represented relative to control cells. (**A**) There is no significant difference (ns) in the luciferase activity after transfection of *Eya1* relative of control or after expression of *Eya1* with *Pa2G4* or Mcrs1 or *Sobp* relative to *Eya1* alone (ns). (**B**) There is no significant difference (ns) in the luciferase activity after transfection of *Eya2* relative of control or after expression of *Eya2* with *Pa2G4* or Mcrs1 or *Sobp* relative to *Eya2* alone (ns).
